# A modular strategy for extracellular vesicle-mediated CRISPR-Cas9 delivery through aptamer-based loading and UV-activated cargo release

**DOI:** 10.1101/2024.05.24.595612

**Authors:** Omnia M. Elsharkasy, Charlotte V. Hegeman, Ivana Lansweers, Olaf L. Cotugno, Ingmar Y. de Groot, Zoë E.M.N.J. de Wit, Xiuming Liang, Antonio Garcia-Guerra, Niels J.A. Moorman, Juliet Lefferts, Willemijn S. de Voogt, Jerney J. Gitz-Francois, Annet C.W. van Wesel, Samir El Andaloussi, Raymond M. Schiffelers, Sander A.A. Kooijmans, Enrico Mastrobattista, Pieter Vader, Olivier G. de Jong

## Abstract

CRISPR-Cas9 gene editing technology offers the potential to permanently repair genes containing pathological mutations. However, efficient intracellular delivery of the Cas9 ribonucleoprotein complex remains one of the major hurdles in its therapeutic application. Extracellular vesicles (EVs) are biological nanosized membrane vesicles released by cells, that play an important role in intercellular communication. Due to their innate capability of intercellular transfer of proteins, RNA, and various other biological cargos, EVs have emerged as a novel promising strategy for the delivery of macromolecular biotherapeutics, including CRISPR-Cas9 ribonucleoproteins. Here, we present a versatile, modular strategy for the loading and delivery of Cas9. We leverage the high affinity binding of MS2 coat proteins (MCPs) fused to EV-enriched proteins to MS2 aptamers incorporated into single guide RNAs (sgRNAs), in combination with a UV-activated photocleavable linker domain, PhoCl. Combined with the Vesicular stomatitis virus G (VSV-G) protein this modular platform enables efficient loading and subsequent delivery of the Cas9 ribonucleoprotein complex, which shows critical dependence on the incorporation and activation of the photocleavable linker domain. As this approach does not require any direct fusion of Cas9 to EV-enriched proteins, we demonstrate that Cas9 can readily be exchanged for other variants, including transcriptional activator dCas9-VPR and adenine base editor ABE8e, as confirmed by various sensitive fluorescent reporter assays. Taken together, we describe a robust and modular strategy for successful Cas9 delivery, which can be applied for CRISPR-Cas9-based genetic engineering as well as transcriptional regulation, underlining the potential of EV-mediated strategies for the treatment of genetic diseases.

## Introduction

The CRISPR (Clustered Regularly Interspaced Short Palindromic Repeats)-Cas9 gene editing system offers the potential to permanently modify or repair genes that contain pathological mutations^1^. As such, CRISPR-Cas9 has significant therapeutic potential in the treatment of a plethora of pathologies with underlying genetic causes. The CRISPR-Cas9 system is based on the use of the prokaryotic Cas9 endonuclease, which is guided to a specific genetic sequence by a single guide RNA (sgRNA)^2,3^. This sgRNA is an engineered RNA fusion molecule, consisting of 2 prokaryotic RNA molecules that are originally expressed as part of the CRISPR-Cas system: a CRISPR RNA (crRNA) and a trans-activating crRNA (tracrRNA)^2^. The crRNA contains a 17-20 nucleotide (nt) targeting sequence, called a spacer sequence, that can bind to a complementary genomic sequence through Watson-Crick base pairing. The tracrRNA forms a complex with the crRNA and serves as scaffold that allows Cas9 to bind the sgRNA. Together, Cas9 and the sgRNA form a ribonucleoprotein (RNP) complex that can generate a double-stranded DNA break at a specific genomic locus that is targeted by the spacer sequence in the sgRNA^4^.

These double-stranded DNA breaks can be repaired by two cellular repair mechanisms: non-homologous end joining (NHEJ) and homology-directed repair (HDR)^5^. In NJEH-mediated repair, both ends of the double-stranded break are directly ligated back to one another. This repair mechanism is relatively error-prone, and may result in insertion or deletion of nucleotides (indels)^6^. If an indel occurs in the coding sequence of a gene, this may lead to frameshift mutations that can result in the truncation and subsequent loss of function of its respective protein. HDR-mediated repair is far less error-prone, as it makes use of a DNA template containing homologous sequences to repair a double-stranded DNA break^7^. Classically, the CRISPR-Cas9 system is employed to knock-out genes by generating NHEJ-mediated indels in target genes, or to repair mutations by co-delivering a DNA template that contains a corrective DNA sequence near the target site^8^.

However, the last decade has seen a vast expansion of additional CRISPR-Cas9 based technologies, by fusing various functional domains to a catalytically inactivated Cas9 protein (dCas9)^9^. Such functional domains may provide a variety of functionalities, such as transcriptional activators or inhibitors which facilitate temporary inhibition or induction of transcription of target genes^10^. Alternatively, Cas9 nickases (nCas9) may be fused to base editing domains to change specific nucleotides within the sgRNA target sequence. Here, an adenine base editor consists of an N-terminal fusion of an adenine deaminase (TadA) to nCas9 resulting in A>G conversion, or a cytosine base editor contains a cytidine deaminase resulting in a C>T conversion^11,12^. These base editors allow for the generation of specific mutations, without the need for an HDR DNA template.

Whereas the CRISPR-Cas9 toolbox has been extensively characterized and expanded over the last decade, intracellular delivery remains one of the major hurdles for therapeutic applications. Efficient intracellular delivery of the Cas9 RNP complex is hampered due to its large size, negative charge, and immunogenicity^13^. To this end, various intracellular delivery strategies have been employed to facilitate Cas9 RNP delivery, including viral vectors, lipid nanoparticles (LNPs), polymers, and cell penetrating peptides (CPPs)^14–17^. Whereas these vectors have shown promising potential, they have substantial limitations. Viral vectors have limited cargo capacity, have increased risk of potential immunogenic responses, and long-term expression of the CRISPR-Cas9 system from viral expression vectors increases the chance of unintended off-target effects, and clearance of target cells by the immune system^18,19^. Both LNPs and polymer-based systems have shown potential for delivery of Cas9 RNP and mRNA, but show cellular toxicity and limited delivery efficiency due to endosomal entrapment^20^. CPPs show similar limitations, and may suffer from decreased particle stability in serum-rich environments^21^. Thus, there is a need for a for a CRISPR-Cas9 delivery strategy that shows low toxicity and immunogenicity, and high stability in serum-rich conditions.

Extracellular vesicles (EVs) are a heterogeneous population of lipid bilayer nanoparticles released by all cell types, and play an important role in intercellular communication^22^. EVs are commonly classified into two major subtypes: exosomes and microvesicles (also referred to as ectosomes). Exosomes range in size from 30 – 200 nm, and are formed in the endosomal pathway through the formation of intraluminal vesicles in the late endosome, resulting in the formation of multivesicular bodies (MVBs)^23^. Upon fusion of MVBs with the cellular membrane, the intraluminal vesicles are released, whereafter they are referred to as exosomes. Microvesicles, or ectosomes, bud directly from the cell membrane and range in size from 100 – 1000 nm^23^. EVs facilitate intercellular communication through transfer of various biological cargos including (trans)membrane proteins, various types of RNA molecules, lipids, and metabolites. Intercellular communication is shown to be facilitated through direct receptor-ligand interactions at the cell surface, or through intracellular delivery of the intraluminal EV cargo in the recipient cell^24^. This form of intercellular communication has been shown to play an important role in a large number of physiological and pathological processes, including immune regulation, tumor progression, angiogenesis, and wound healing^25–28^.

Given their innate capacity of intercellular transfer of biological cargos, capability of crossing difficult-to-cross biological barriers, and their low immunogenicity, EVs have raised increasing interest as potential vectors for therapeutic delivery of a variety of biological cargos including proteins and RNA. Various studies have shown that these natural nanoparticles can be enriched with therapeutic proteins during biogenesis by expressing fusion proteins of EV-enriched proteins and the protein of interest in the EV-producing cells^29,30^. Such EV-enriched proteins commonly include tetraspanins CD9, CD63, and CD81. One potential limitation of this approach for Cas9 loading, is that fusion of an EV-enriched protein directly to Cas9 may interfere with the functionality of the aforementioned enzymatic domains present on the C-terminus or the N-terminus of Cas9. Moreover, as Cas9 needs to translocate to the nucleus upon delivery, tethering of Cas9 to the EV membrane may prevent its transfer to the appropriate cellular compartments in recipient cells. Thus, in order to generate a modular loading strategy that can be applied to load and deliver a variety of Cas9-based technologies through EVs, alternative EV loading strategies are preferred.

Here, we present a modular and versatile strategy to facilitate EV-mediated delivery of various Cas9-based applications. This strategy is based on the fusion of RNA-binding domains to EV-enriched moieties, separated by a UV-cleavable linker^31,32^. The RNA-binding domains within these fusion proteins facilitate the loading of Cas9 RNPs into EVs by high-affinity binding of aptamers that are incorporated into the tetraloop and second stem loop of the Cas9 sgRNA^33^. As such, no direct fusion to either the C- or N-terminus of Cas9 is required, providing a flexible platform for delivery of a multitude of Cas9 variants. As cleavage of the UV-cleavable linker can be efficiently activated after EV isolation, this approach shows efficient delivery of various Cas9 variants, including transcriptional activation and adenine base editing, using multiple reporter systems. Altogether, this work demonstrates the development of a versatile and efficient EV-based Cas9 RNP delivery platform.

## Materials & Methods

### Cell culture

HEK293T cells (CRL-3216) and MDA-MB-231 cells (HTB-26) were obtained from the American Type Culture Collection (ATCC), and were both cultured in Dulbecco’s Modified Eagle Medium (DMEM) with L-Glutamine (Gibco) supplemented with 10% fetal bovine serum (FBS) (Sigma-Aldrich). HEK293T stoplight cells and HEK293T adenine base editor stoplight cells were generated as previously described^17,34^. HEK293T cells with stable doxycycline-inducible eGFP expression were generated for this study as described below. All cell lines were cultured at 37 °C and 5% CO_2_.

### DNA constructs

For unmodified sgRNA expression, targeting sequences were cloned into a lentiGuide-Puro plasmid (Addgene #52963)^35^ as previously described^34^. In short, complementary synthesized oligonucleotides (Integrated DNA technologies) were annealed and ligated into a lentiGuide-Puro plasmid after BsmBI digestion (New England Biolabs). For the expression of MS2-sgRNAs, complementary synthesized oligonucleotides were annealed and cloned into the lenti sgRNA(MS2)-Zeo backbone (Addgene #61427)^33^ after BsmBI digestion. For expression of sgRNAs with a single MS2 hairpin integration, sgRNA sequences were synthesized as gBlocks (Integrated DNA technologies), and cloned into a Lenti_gRNA-Puro backbone (Addgene #84752)^36^ after BsmBI digestion. Oligonucleotide and gBlock sequences used for cloning sgRNA expression constructs are listed in Supplementary Table 1, expressed sgRNA sequences are listed in Supplementary Table 2. For the generation of a lentiviral doxycycline-inducible eGFP expression construct, an eGFP open reading frame was transferred from pDONR221- eGFP (Addgene #25899)^37^ into pInducer20 (Addgene #44012)^38^ using the Gateway LR Clonase II Enzyme Mix (Thermo Fisher Scientific) according to the manufacturer’s protocol. For expression of MCP loading constructs, including those with photocleavable PhoCl domains, human codon-optimized sequences were synthesized as gBlocks, and cloned into pHAGE2- EF1a-Multiple Cloning Site-IRES-PuroR plasmids using NotI and BamHI restriction enzymes (New England Biolabs), as described previously^39^. Amino acid sequences for all RNA-binding constructs are shown in Supplementary Table 3. An overview of used plasmid DNA constructs is listed in Supplementary Table 4.

### Lentiviral production and generation of stable cell lines

For lentiviral production HEK293T cells were plated at 50% confluency in DMEM supplemented with 10% FBS and supplemented with 1x Antibiotic Antimycotic Solution (Sigma-Aldrich), and transfected with lentiviral transfer plasmids containing genes of interest (GOI), PSPAX2 (Addgene #12260) and pMD2.G (Addgene #12259) at a 2:1:1 ratio using 3 μg 25 kDa linear polyethylenimine (PEI) (Polysciences Inc.) per μg DNA. After 18 hours culture medium was replaced with fresh DMEM supplemented with 10% FBS and 1x Antibiotic Antimycotic Solution. After 48 hours, lentiviral supernatants were harvested. Cells were removed by a 10 minute centrifugation at 500 x *g*, followed by 0.45 μm syringe-filtration (Sartorius). Cells were transduced overnight using lentiviral stocks supplemented with 8 μg/ml polybrene (Sigma-Aldrich), after which lentiviral supernatant was replaced by fresh culture medium. 24 hours after lentiviral transduction, cells were cultured with their respective selection antibiotics. All reporter cell lines were cultured in the presence of 1000 μg/ml G418 (Invivogen). Stable donor cell lines expressing MS2-sgRNA were cultured in the presence of 200 μg/ml Zeocin (Invivogen), and stable donor cell lines expressing MCP-CD63 were cultured in the presence of 2 μg/ml puromycin (Invivogen).

### Plasmid transfection

For transfection of plasmid DNA for expression of EV cargos, HEK293T cells were plated at 1.0 x 10^7^ cells per T175 flasks in culture medium supplemented with 1x Antibiotic Antimycotic Solution. After 24 hours, HEK293T cells were transfected with 5 μg for each plasmid per flask. All plasmids were mixed in a 50 ml conical tube in a volume of 1 ml OptiMEM per transfected flask, and simultaneously 2 μg 25 kDa linear PEI per μg plasmid DNA was mixed in a separate conical tube in a volume of 1 ml OptiMEM per transfected flask. After a 5 minute incubation at room temperature, the contents of both tubes were transferred to a single conical tube, and gently mixed by inverting the tube several times. After a 15 minute incubation at room temperature, 2 ml transfection mixture was added to each T175 flask, followed by an overnight transfection at 37 °C and 5% CO_2_. Then, the transfection medium was removed and the cells were gently washed using 5 ml OptiMEM per flask. Hereafter, 20 ml OptiMEM supplemented with 1x Antibiotic Antimycotic Solution was added per flask. After 24 hours, conditioned medium was isolated for subsequent EV isolation as described below. For direct transfection of HEK293T cells to study reporter cell activation, cells were plated in 24-well plate wells at a density of 5.0 x 10^4^ cells per well in 1 ml culture medium supplemented with 1x Antibiotic Antimycotic Solution, 24 hours priors to transfection. HEK293T reporter cells were transfected with 250 ng per plasmid per well, unless stated otherwise. All plasmids were mixed in a 1.5 ml tube in a volume of 50 μl OptiMEM per transfected well, and simultaneously 3 μg 25 kDa linear PEI per μg plasmid DNA was mixed in a separate 1.5 ml tube in a volume of 50 μl OptiMEM per transfected well. After a 5 minute incubation at room temperature, both solutions were transferred to a single 1.5 ml tube and gently mixed. After a 15 minute incubation at room temperature, 100 μl transfection mixture was added to each well. Reporter cells were then analyzed by flow cytometry and fluorescence microscopy as described below. Reporter cells were analyzed at least 48 hours after transfection, to ensure sufficient levels of eGFP expression.

### Extracellular vesicle isolation

HEK293T cells were plated in T175 culture flasks for EV isolation in DMEM supplemented with 10% FBS. HEK293T were plated to be at 80% confluency 24 hours prior to EV isolation. Cell culture medium was removed, and the cells were gently washed with 5 ml OptiMEM per flask. Hereafter, 20 ml OptiMEM supplemented with 1x Antibiotic Antimycotic Solution was added per flask. After 24 hours, conditioned medium was isolated for extracellular vesicle isolation. First, cells and cellular debris were removed by a 5 minute centrifugation step at 300 x *g*, followed by a 15 minute centrifugation step at 2,000 x *g*. Supernatant was then collected and filtered through a 0.45 μm vacuum filter (Corning). The filtered supernatant was then concentrated by tangential flow filtration (TFF), using a Vivaflow 50R 100 kDa TFF cassette (Sartorius) to a volume of 15 ml, and subsequently concentrated to a 0.5 – 1.0 ml volume using 100 kDa Amicon Ultra-15 Centrifugal filters (Merck). EVs were then isolated by size exclusion chromatography (SEC) on an Akta Pure chromatography system using a Tricorn 10/300 column with Sepharose 4 Fast Flow resin (all GE Healthcare Life Sciences). EVs were then sterilized by 0.45 μm syringe-filtration, and concentrated to a 300 – 400 μl volume using 10 kDa Amicon Ultra-15 Centrifugal filters (Merck).

### Nanoparticle tracking analysis

EV particle concentration and size distribution was determined using a Nanosight S500 nanoparticle analyzer, equipped with a 405 nm laser (Malvern Instruments). Samples were diluted in PBS (Sigma-Aldrich) to a suitable concentration for nanoparticle tracking analysis (between 1.5 x 10^8^ and 1.5 x 10^9^ particles per ml) and were analyzed using 5 x 30 second recordings per sample with a camera sensitivity setting at level 16. All acquisition settings were set to “Auto” for post-acquisition analysis, with the exception of a fixed detection threshold, which was set to 7. Recordings were analyzed using NTA software v3.4. Particle counts were corrected for measurements of PBS, followed by correction for dilution.

### Western blot

Cells or isolated EVs were lysed in RIPA buffer, supplemented with a protease inhibitor cocktail (Sigma-Aldrich). Cell lysates were incubated on ice for 30 minutes, followed by a 15 minute 4 °C centrifugation step at 12,000 x *g* to remove non-soluble materials. Protein concentrations were measured using a BCA Protein Assay Kit (Thermo Fisher Scientific), alongside a BSA protein standard (Thermo Fisher Scientific), according to the manufacturer’s protocol. Samples were normalized for protein concentration, and mixed with premixed sample loading buffer (Bio-Rad). With the exception of CD63 western blot analysis, 100 µM DTT reducing agent was included in the sample loading buffer. Prior to western blot analysis, samples were denatured by a 10 minute incubation at 95 °C. Then, samples were loaded on 4–12% gradient Bis–Tris polyacrylamide gels in 1x MOPS buffer (Thermo Fisher Scientific), and subjected to electrophoresis. Hereafter, proteins were blotted onto Immobilon-FL PVDF membranes (Millipore), which was subsequently blocked for 1 hour at room temperature using a blocking buffer consisting of 1 part Intercept Blocking Buffer (LI-COR Biosciences) and 1 part Tris-Buffered Saline (TBS). Membranes were then probed overnight at 4 °C in staining buffer consisting of 1 part Intercept Blocking Buffer (LI-COR Biosciences) and 1 part Tris-Buffered Saline with 0.1% Tween-20 (TBS-T), using the following primary antibodies: ALIX 1:1000 (Thermo Fisher Scientific, MA1-83977), Calnexin 1:1000 (GeneTex, GTX101676), CD63 1:1000 (AB8219), TSG101 1:1000 (Abcam, ab30871), and H2B 1:1000 (Abcam, ab52599), GAPDH 1:500 (Abcam, AB9485) or α-Flag 1:1000 (Sigma-Aldrich, F1804). Blots were washed 5 times for 5 minutes at room temperature in TBS-T, followed by a 2 hour probe with secondary antibodies in staining solution at room temperature, using the following secondary antibodies at a 1:10,000 dilution: anti-rabbit IgG conjugated to AlexaFluor 680 (Thermo Fisher Scientific, A-21076), anti-mouse IgG conjugated to AlexaFluor 680 (Thermo Fisher Scientic, A- 21057), anti-rabbit IgG conjugated to IRDye 800CW (926-32211, LI-COR Biosciences), or anti-mouse IgG conjugated to IRDye 800CW (926-32212, LI-COR Biosciences). Next, blots were washed 3 x 5 minutes at room temperature in TBS-T, and 2 x 5 minute washing steps at room temperature in TBS. Fluorescent imaging was done using an Odyssey Infrared Imager (LI-COR Biosciences) at 700 and 800 nm.

### Transmission electron microscopy

EVs were isolated as described above, and were subsequently adsorbed to carbon-coated coated formvar grids (TAAB Laboratories Equipment Ltd.) at room temperature for 15 minutes. Formvar grids were washed with PBS to remove any remaining unbound EVs, and grids were subsequently fixed using a fixing buffer (2% paraformaldehyde, 0.2% glutaraldehyde in PBS) for 30 minutes at room temperature. After counterstaining with uranyl-oxalate, grids were embedded an 1.8% methyl cellulose and 0.4% uranyl acetate mixture at 4 °C. Grids were imaged on a Jeol JEM-1011 microscope (Jeol).

### Optiprep density gradient

After EVs were isolated as described above, EV SEC fractions were transferred to SW40 centrifuge tubes (Beckman Coulter), PBS was added to a volume of 14 ml, and EVs were pelleted by ultracentrifugation for 60 minutes at 100,000 x *g* at 4 ° C in an SW40.Ti rotor (Beckman Coulter). After centrifugation, the EV pellet was resuspended in 3 ml 40% Optiprep (Sigma-Aldrich) in PBS. Hereafter, 3 ml fractions of 30%, 20%, and 10% Optiprep were gently layered on top, and centrifuged for 16 hours at 4° C at 200,000 x *g* in an SW40.Ti rotor. 1 ml fractions were then gently isolated from the top of the Optiprep gradient and transferred to 2 ml tubes. Next, proteins were precipitated for each of the isolated fractions using trichloroacetic acid (TCA) precipitation. 250 µl TCA was added to each 1 ml fraction and mixed thoroughly. After an overnight precipitation at −20 °C, proteins were pelleted by centrifugation at 10,000 x *g* for 10 minutes at 4 °C. After supernatant removal, pellets were washed 3 times with 500 µl ice-cold acetone, followed by centrifugation for 10 minutes at 10,000 x *g* at 4 °C. After the final washing step supernatant was removed, and pellets were dried at 95 °C for 10 minutes. Samples were then resuspended in sample buffer and analyzed by western blot analysis as described above.

### RNA isolation

To assess EV sgRNA loading by qPCR without interference of plasmid DNA transfection complexes, stable MS2-sgRNA expressing MDA-MB-231 cell lines with- or without stable co-expressing of MCP-CD63 were generated as described above. RNA was isolated from cells using TRIzol reagent (Thermo Fisher Scientific), and from EVs using TRIzol LS reagent (Thermo Fisher Scientific), according to manufacturer’s protocol using glycoblue as co-precipitant (Thermo Fisher Scientific). RNA pellets were resuspended in nuclease-free water (Thermo Fisher Scientific), and RNA concentration and purity were determined using a Nanodrop N1000 (Thermo Fisher Scientific).

### RT-PCR and qPCR analysis

For RT-PCR analysis, 1 μg of RNA was used for reverse transcription using an iScript cDNA synthesis kit (Bio-Rad). cDNA was diluted 5 times in Milli-Q water and used as a template for PCR amplification using GoTaq G2 Green Master Mix (Promega) according to the manufacturer’s protocol. PCR products were loaded directly onto a 2% TAE agarose gel with GelRed Nucleic Acid Gel Stain (Sigma-Aldrich) for electrophoresis, and imaged using a Bio-Rad Gel DOC XR+ imager (Bio-Rad). For qPCR analysis, 500 ng RNA was used for reverse transcription using the SuperScript IV cDNA synthesis kit (Thermo Fisher Scientific). cDNA was diluted 5 times in Milli-Q water, whereafter 5 μl was used for a 25 μl qPCR reaction using 2x SYBR Green Master Mix (Bio-Rad) in a CFX96 Real-Time PCR Detection System (Bio-Rad) according to the manufacturer’s protocol. Cycle threshold (Ct) values were corrected for the housekeeping gene GAPDH. PCR primers were synthesized by Integrated DNA Technologies, and sequences are listed in Supplementary Table 5.

### Extracellular vesicle addition experiments

To study EV-mediated Cas9 RNP delivery, EVs were added to cells in 24-well plate wells, cultured in 1 ml culture medium. Unless stated otherwise, 5.0 x 10^10^ EVs were added per 24- well plate well containing 1.0 x 10^5^ reporter cells. EV dosages were normalized based on nanoparticle tracking analysis, as described above. Prior to EV addition, EVs containing loading constructs with PhoCl domains were treated with UV light on ice for 20 minutes, using a custom 50W 395 nm LED panel (SJLA). Reporter cells were analyzed 72 hours after EV addition to ensure sufficient levels of eGFP expression, unless stated otherwise. Reporter cells were analyzed by flow cytometry and fluorescence microscopy.

### Fluorescence microscopy

Prior to flow cytometry analysis, reporter cells were analyzed by fluorescence microscopy using either an EVOS FL Cell Imaging System (Thermo Fisher Scientific), or a Nikon Eclipse TS2 Inverted Fluorescence Microscope (Nikon Instruments). Images were processed using FIJI ImageJ v1.54d software.

### Flow cytometry

After fluorescence microscopy, cells were gently washed with 0.5 ml PBS and trypsinized for 5 minutes using 250 μl TrypLE Express (Thermo Fisher Scientific). Cells were then transferred to 1.5 ml tubes using a similar volume of cell culture medium containing 10% FBS. Cells were centrifuged for 5 minutes at 300 x *g*, washed in 1 ml 1% FBS in PBS, and centrifuged for 5 minutes at 300 x *g*. After supernatant removal, cells were resuspended in 200 μl 1% FBS in PBS, transferred to 96-well U-bottom Falcon plates (Fisher Scientific), and analyzed on a BD FACSCanto II flow cytometry system (BD Biosciences), or a BD LSRFortessa Cell Analyzer (BD Biosciences). Flow cytometry results were analyzed using FlowJo v10 software. Gating strategies for all reporter cell lines are shown in Supplementary Figure 1.

### Statistics

All statistical analyses were performed using GraphPad Prims v10.2 software. Unless stated otherwise, values are presented as mean ± standard deviation. Two-sided statistical tests were performed in all statistical analyses within this manuscript. Statistical significance was considered at *p* < 0.05.

## Results

### Generation of a modular aptamer-based strategy Cas9-sgRNA RNP loading into EVs

As the interest in Cas9-mediated gene therapy has increased over the last decade, the abundance of Cas9-based tools, such as transcriptional regulators and base editors, has grown along with it. In order to generate a universal loading strategy that can be applied interchangeably to all these variants, we opted to avoid the use of direct Cas9-tetraspanin fusion proteins. Instead, we intraluminally fused tandem MS2 coat proteins (MCPs), lacking the Fg loop to prevent capsid formation, to the N-terminus of the EV-enriched tetraspanin CD63 (Fig. 1A). These MCP proteins robustly bind to MS2 aptamers, specific RNA hairpin sequences. To bind the Cas9-sgRNA ribonucleoprotein (RNP) complex, MS2 aptamers that can be bound by the tandem MCPs were placed in the tetraloop and second stemloop of the sgRNA. Previous studies have shown that replacing the hairpins in these sections of the sgRNA does not interfere with the Cas9 RNP functionality, and that these hairpins protrude from the Cas9 RNP structure^33^. In this modular approach, sgRNAs and Cas9 are readily interchangeable, as no direct fusion proteins are employed.

**Figure 1.**
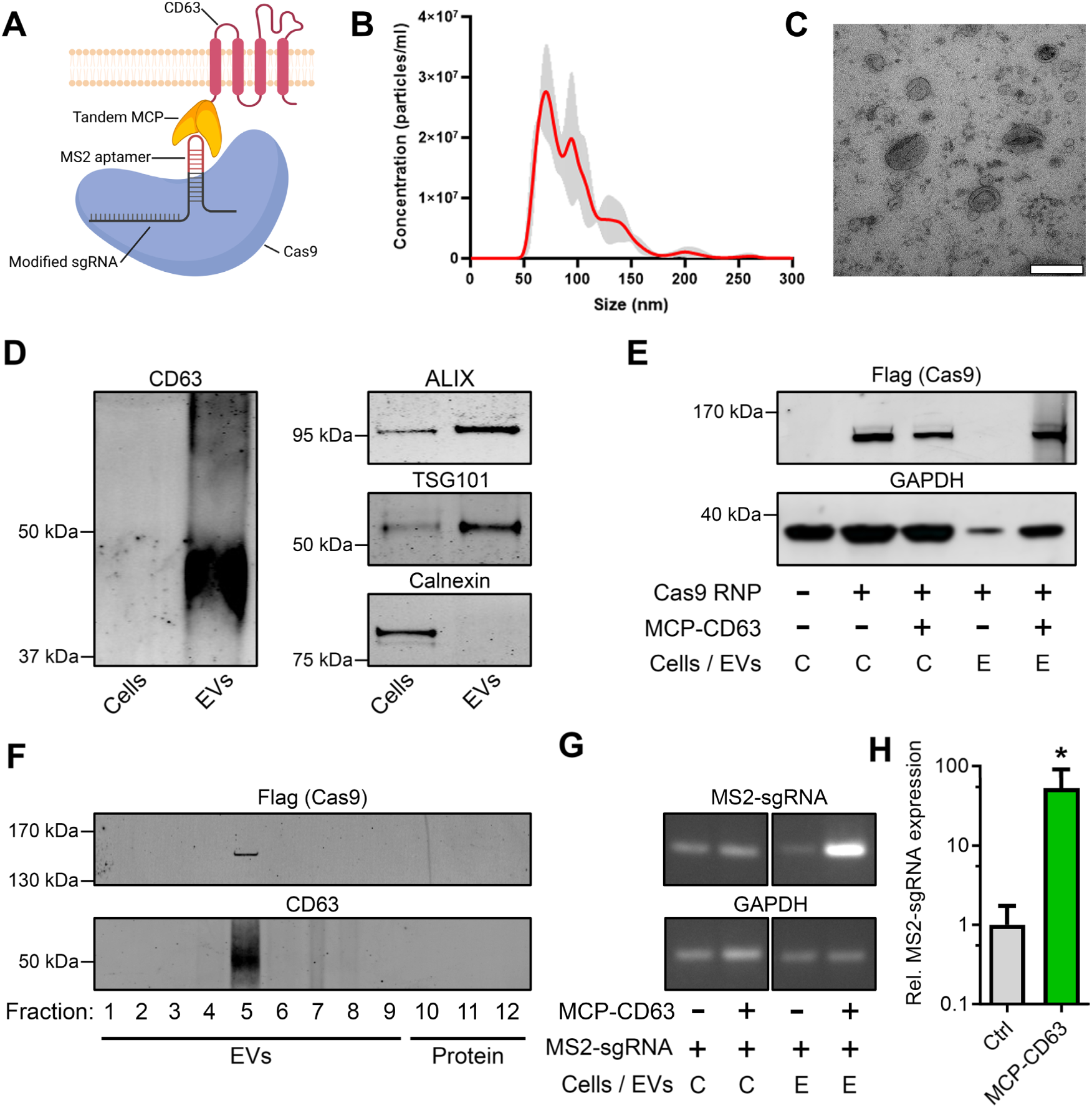
Engineering EVs for the targeted loading of Cas9 ribonucleoprotein complexes. **a,** Schematic of the EV engineering strategy for active loading and delivery of Cas9 RNPs in HEK293T-derived EVs. Tandem MCPs, lacking the Fg loop involved in capsid formation, are intraluminally fused to the N-terminus of EV-enriched CD63 (MCP-CD63). These MCPs bind MS2 aptamers protruding from the RNP, which are present on MS2-modified sgRNAs. **b,** Nanosight particle tracking analysis displaying the size distribution of isolated EVs. **c,** Electron microscopy image of isolated EVs, scalebar represents 200 nm. **d,** Western blot analysis of cell lysates and EVs shows an enrichment for EV markers CD63, ALIX and TSG101, and a negative enrichment for the ER organelle marker Calnexin, in isolated EVs. Due to the highly glycosylated nature of CD63, its western blot analysis presents a commonly observed “smear” pattern. **e,** Western blot analysis shows high levels of Cas9 in cell lysates regardless of co-expression of MCP-CD63, whereas high levels of Cas9 in isolated EVs is only observed upon co- expression of MCP-CD63. **f,** Western blot analysis of an OptiPrep density gradient of isolated EVs shows presence of Cas9 in the same EV-associated fractions as EV-marker CD63. **g, h** RT-PCR analysis **(g)** of RNA isolated from cells and EVs from stably expressing MS2-sgRNA with- or without the stable co-expression of MCP-CD63, and qPCR-analysis of RNA isolated from EVs **(h)**, shows enriched loading of MS2-sgRNAs into isolated EVs. Means + SD, n = 5, Student’s *t*-test. * *p* < 0.05.

HEK293T cells were transfected with plasmids expressing the MCP-CD63 loading construct, Cas9, and MS2-sgRNAs, and 48 hours later EVs were isolated by tangential flow filtration (TFF), followed by size exclusion chromatography (SEC). Nanoparticle tracking analysis (NTA, Fig. 1B) and transmission electron microscopy (TEM, Fig. 1C) confirmed the isolation of particles with a morphology and size distribution (mode = 75.1 +/- 5.1 nm) in line with the common observed characteristics of EVs. Additionally, western blot analysis showed enrichment for common EV-markers CD63, ALIX and TSG101, and a negative enrichment for the organelle marker Calnexin, as compared to cell lysates. To assess whether expression of our MCP-CD63 loading construct indeed resulted in the targeted loading of Cas9 RNPs into EVs HEK293T cells were transfected with plasmids for expressing Cas9 and MS2-sgRNA, with- or without the co- expression of MCP-CD63. Western blot analysis of EV and cell lysates revealed that Cas9 was present in cell lysates independent of the co-expression of MCP-CD63, but was only found in EVs when MCP-CD63 was co-expressed, confirming the efficacy of this loading strategy (Fig. 1E). To ensure that the observed Cas9 protein was indeed associated with EVs, isolated EVs were further analyzed using an OptiPrep density gradient, wherein EVs can be separated from non-membrane-associated contaminants by density-based separation. Indeed, presence of Cas9 was only observed in EV-associated fractions positive for EV-marker CD63 (Fig. 1F). Similarly, to assess whether MS2-sgRNA was also actively enriched in EVs by MCP-CD63, MS2- sgRNA loading was analyzed by RT-PCR (Fig. 1G) and qPCR (Fig. 1H). As was also observed for Cas9 protein, co-expression of MCP-CD63 resulted in a significant increase in MS2-sgRNA abundance in isolated EVs.

To confirm and analyze EV-mediated Cas9 RNP delivery to target cells in a sensitive and robust manner, we employed a fluorescent reporter for Cas9 activity that we previously published^34^. In this fluorescent “stoplight reporter system”, an mCherry open reading frame (ORF), followed by a short linker region that contains a Cas9 target site, is constitutively expressed. When Cas9 and a targeting sgRNA are functionally delivered, a resulting frameshift can activate one of the out-of-frame downstream eGFP ORFs, resulting in permanent high eGFP expression (Fig. 2A). First, to test whether inclusion of MS2 aptamers in the sgRNA tetraloop and second stemloop indeed did not interfere with RNP functionality, HEK293T cells expressing the stoplight reporter construct were transfected with Cas9 in combination with a non-targeting (NT) sgRNA, a targeting (T) sgRNA, or a targeting MS2 (T MS2) sgRNA, and analyzed by fluorescence microscopy (Fig. 2B) and flow cytometry (Fig. 2C, Fig. S1A). Both analyses showed no activation of eGFP expression when Cas9 was co-transfected with an NT sgRNA, and showed high levels of activation when Cas9 was co-transfected with both the wildtype T sgRNA and the T MS2-sgRNA. Moreover, no significant decrease in Cas9-mediated activation of eGFP expression was observed in T MS2-sgRNA as compared to WT T sgRNA, confirming that this modification does not interfere with Cas9 activity. Next, the capacity of our engineered EVs to deliver Cas9 RNPs was assessed using the stoplight reporter construct. To further enhance EV-mediated Cas9 delivery, EV-producing cells were also transfected to express the Vesicular stomatitis virus (VSV) envelope glycoprotein VSV-G^40^. This glycoprotein facilitates endosomal escape of delivered cargo in recipient cells by inducing membrane fusion in the late endosome, triggered by the local decrease in pH. To test the effect of MCP- CD63 on EV-mediated Cas9 delivery, EVs from cells expressing VSV-G, Cas9 and MS2-sgRNA, with- or without MCP-CD63 co-expression, were isolated and added to HEK293T stoplight reporter cells. Whereas both flow cytometry (Fig. 2D) and fluorescence microscopy (Fig. 2E) confirmed an increase in EV-mediated Cas9 delivery due to MCP-CD63 expression, levels of recombination in the reporter cells were surprisingly low, at less than 2%.

**Figure 2.**
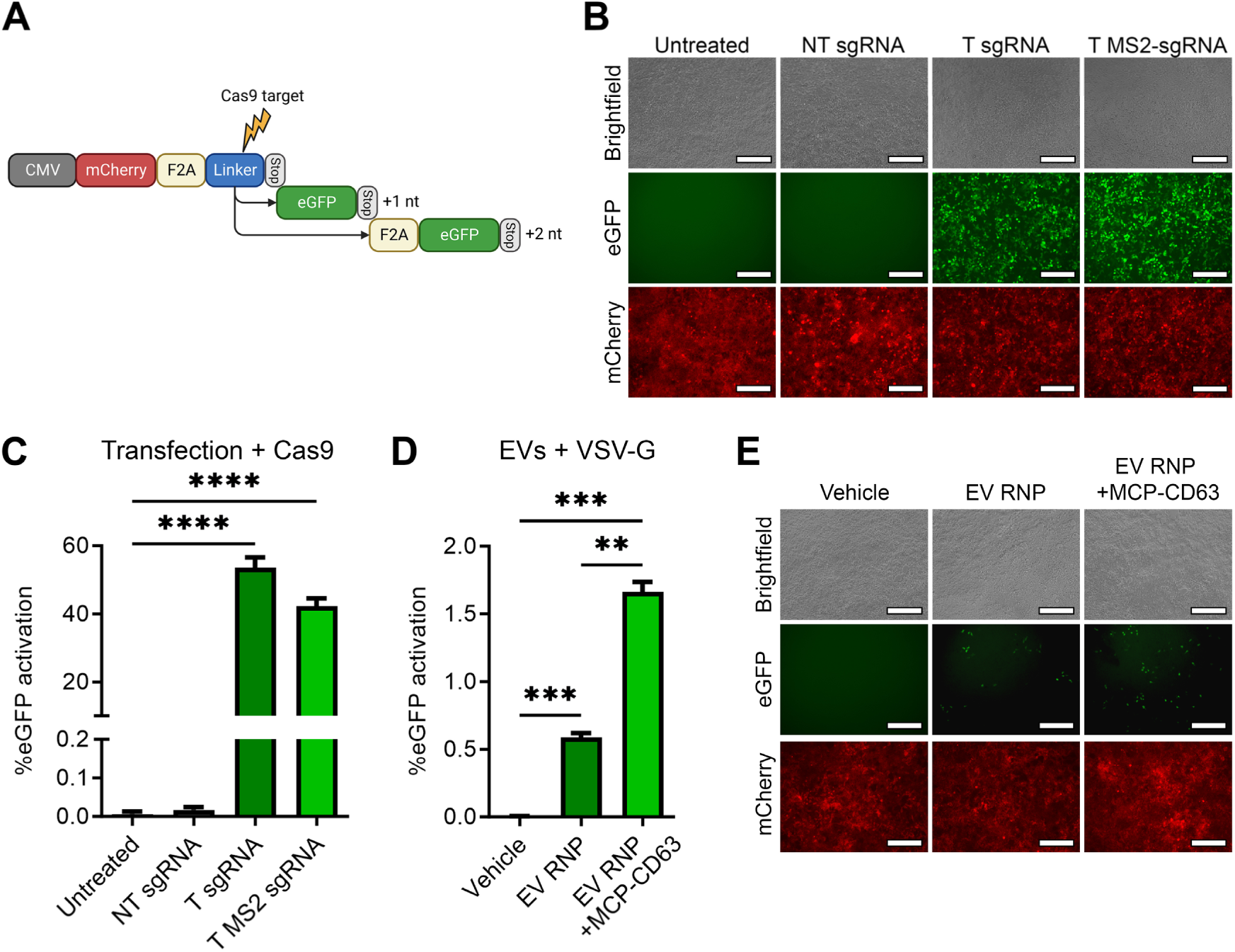
MCP-CD63 facilitates a limited increase of EV-mediated Cas9 RNP delivery. **a,** Schematic of the fluorescence “stoplight” reporter construct for Cas9 activity. mCherry (red) is stably expressed under a CMV promoter followed by a small “linker” region (blue), containing a Cas9 target site, and a stop codon. Frameshifts in the linker region, resulting from non-homologous end joining (NHEJ)-mediated repair mechanisms, due to Cas9-mediated double stranded breaks result in expression of downstream eGFP (green) open reading frames. **b, c** Fluorescence microscopy images **(b)** and flow cytometry analysis **(c)** of HEK293T cells expressing the stoplight reporter construct, transfected with Cas9 and a non-targeting (NT) sgRNA, or a targeting (T) sgRNA with or without MS2 aptamers. eGFP expression is observed after transfection of Cas9 with both wildtype targeting (T) sgRNA or a T MS2-sgRNA. Scalebar represents 200 μm. Means + SD, n = 3, Dunnett’s multiple comparison test. **d, e** Flow cytometry analysis **(d)** and fluorescence microscopy images **(e)** of HEK293T cells expressing the stoplight reporter construct, 72 hours after addition of EVs isolated from HEK293T cells expressing Cas9 + MS2- sgRNA + VSV-G (EV RNP) or expressing Cas9 + MS2-sgRNA + MCP-CD63 + VSV-G (EV RNP + MCP-CD63) shows that MCP-CD63 facilitates a significant but limited increase of EV-mediated RNP delivery. Scalebar represents 200 μm. Means + SD, n=3, One-way ANOVA with post-hoc Tukey’s multiple comparisons test. ** *p* < 0.01, *** *p* < 0.001, **** *p* < 0.0001.

### Incorporation of a photocleavable domain facilitates efficient cargo release and delivery

Since the significant increase in MS2-sgRNA and Cas9 loading by MCP-CD63 did not result in a similar level of increase in Cas9 delivery, we hypothesized that the loaded Cas9 RNP was not sufficiently released from the EV membrane due to the strong binding affinity of the MS2 aptamers in the sgRNA to the MCPs tethered to CD63. As a result, Cas9 would be unable to translocate into the nucleus of the target cell. To address this issue, we introduced the PhoCl photocleavable protein between the tandem MCP domains and the CD63 tetraspanin (Fig. 3A)^32^. This PhoCl protein is a derivate of the FP mMaple protein, and is cleaved when exposed to 400 nm (ultra)violet light. To confirm that this construct was indeed cleaved upon UV exposure, HEK293T cells were transfected with the MCP-PhoCl-CD63 construct, and after 48 hours were exposed to UV light from a 50W 395 nm LED panel for various lengths of time. As the PhoCl domain was given an HA-tag, protein cleavage from the ∼82 kDa fusion protein into a ∼55 kDa subunit could be observed by western blot analysis (Fig. 3B). Here, protein cleavage was confirmed, and optimal cleavage was achieved in a 15 – 30 minute timeframe. UV exposure to EVs isolated from cells expressing the MCP-PhoCl-CD63 construct showed similar kinetics, were optimal cleavage was reached at around 20 minutes UV exposure (Fig. 3C). Given these promising observations, EVs from cells expressing VSV-G, Cas9, MS2-sgRNA, and MCP-PhoCl-CD63 were isolated, and subsequently used for direct addition to stoplight reporter cells, or were first exposed to UV light for 20 minutes on ice prior to addition to stoplight cells. 48 hours after EV addition, flow cytometry analysis (Fig. 3D) and fluorescence microscopy (Fig. 3E) both confirmed a significant and substantial increase in EV-mediated Cas9 delivery in MCP-PhoCl-CD63 EVs treated with UV, increasing recombination in reporter cells from ∼2% to ∼28%. To assess the importance of including the VSV-G envelope glycoprotein, EVs from cells expressing Cas9 and MS2-sgRNA, with- or without co-transfection of MCP-CD63 (Fig. S2A) or MCP-PhoCl-CD63 +/- UV treatment (Fig. S2B) without co-expression of VSV-G were added to stoplight reporter cells. In all conditions flow cytometry analysis showed no increase in eGFP expression. These data indicate that, despite the observed high increase in Cas9 RNP loading in presence of MCP-CD63, and the high increase in EV-mediated Cas9 RNP delivery in presence of MCP-PhoCl-CD63 with UV treatment, co-expression of VSV-G is pivotal for efficient Cas9 delivery in this approach. To assess whether the 395 nm UV treatment of the EVs prior to addition to recipient cells did not have adverse effects on the functionality of the loaded Cas9 RNPs, VSV-G+ EVs loaded with MCP-CD63 or MCP-PhoCl- CD63 were used for an addition assay, with- or without UV treatment. As observed before, flow cytometry analysis (Fig. S2C) and fluorescence microscopy (Fig. S2D) showed that UV treatment of MCP-PhoCl-CD63 EVs highly increased Cas9 delivery. Importantly, we also observed that UV treatment of MCP-CD63 EVs did not result in a decrease in Cas9 delivery, confirming that the UV treatment did not negatively affect the EV-associated Cas9 RNP functionality.

**Figure 3.**
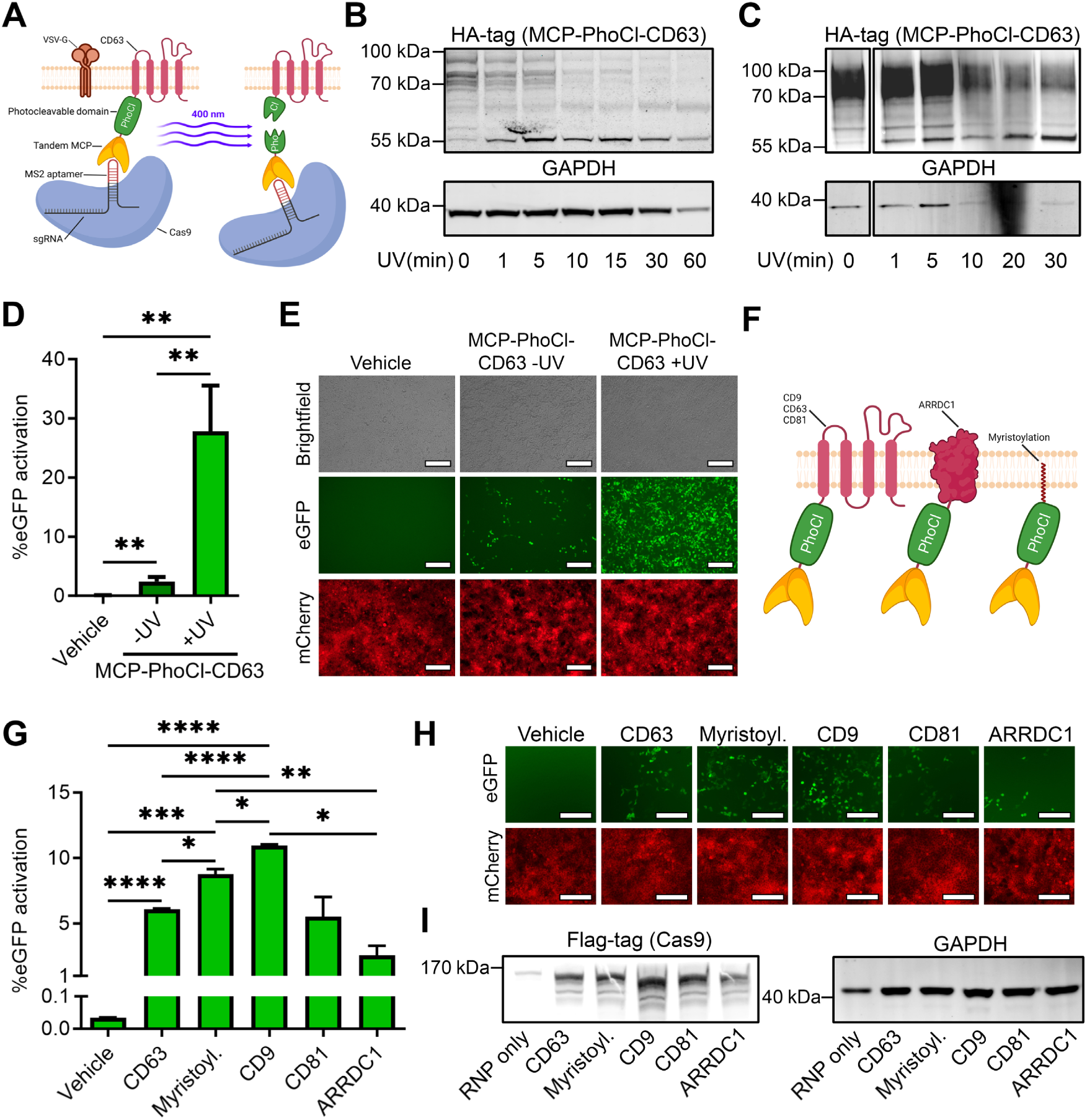
Incorporation of a photocleavable domain to facilitate cargo release strongly increases EV-mediated Cas9 RNP delivery. **a,** Schematic of the EV engineering strategy for photo-activatable release of membrane bound MCP-loaded Cas9 RNPs. A photocleavable domain (PhoCl), with a N-terminal HA-tag for western blot analysis, is placed between the tandem MCPs and CD63 (MCP-PhoCl-CD63). Upon exposure to 400 nm UV light PhoCl is cleaved, releasing the MCP-RNP complex from the EV membrane. **b,c** Western blot analysis of MCP- PhoCl-CD63 cleavage in cells **(b)** and isolated EVs **(c)**. Upon UV exposure, the ∼82 kDa fusion protein is cleaved, revealing a ∼55 kDa cleavage product. **d,e** Flow cytometry analysis **(d)** and fluorescence microscopy images **(e)** of HEK293T cells expressing the stoplight reporter construct, 72 hours after addition of EVs isolated from HEK293T cells expressing Cas9 + MS2-sgRNA + MCP-PhoCl-CD63 + VSV-G shows that UV treatment of EVs prior to addition to cells strongly increases EV-mediated RNP delivery. 1.0x10^12^ EVs per well. Mean + SD, n = 5, One-way ANOVA with post-hoc Tukey’s multiple comparisons test. Scalebar represents 200 μm. **f,** Schematic of additional EV-targeted loading constructs. Tandem MCPs are fused to EV-enriched moieties CD9, CD63, CD81, ARRDC1, and a myristoylation tag via a photocleavable (PhoCl) domain. **g,h** Flow cytometry analysis **(g)** and fluorescence microscopy images **(h)** comparing EV-mediated RNP delivery of various MCP-PhoCl fusion proteins. Addition normalized by particle count; 5.0x10^10^ particles added per well. Scalebar represents 200 μm. Means + SD, n = 3, One-way ANOVA with post-hoc Tukey’s multiple comparison test. **i,** Western Blot analysis of Cas9 loading in EVs by various MCP-PhoCl fusion proteins. GAPDH analysis is included as a loading control. * *p* < 0.05, ** *p* < 0.01, *** *p* < 0.001, **** *p* < 0.0001.

Having established an efficient platform for EV-mediated Cas9 loading and delivery, we opted to test the effect of replacing CD63 with alternative EV-enriched moieties on EV-mediated Cas9 delivery efficiency. To this end, CD63 was replaced with tetraspanins CD9 or CD81, Arrestin Domain Containing 1 (ARRDC1), or an N-terminal myristoylation tag (Fig. 3F). EVs were isolated from cells expressing these loading constructs alongside VSV-G, MS2-sgRNA and Cas9, normalized by nanoparticle tracking analysis, treated with 395 nm UV, and added to stoplight reporter cells. Over the course of multiple experiments, flow cytometry analysis (Fig. 3G) and fluorescence microscopy (Fig. 3H) both confirmed that the choice of EV-enriched fusion protein has a significant effect on EV-mediated Cas9 delivery levels. Whereas CD81 performed similarly to CD63, both CD9 and the myristoylation tag showed significantly higher levels of Cas9 delivery, with CD9 outperforming CD63 almost 2-fold. Interestingly, the ARRDC1-mediated microvesicle (ARMM) marker ARRDC1 showed substantially lower levels of Cas9 delivery, despite its previously reported use for efficient Cas9 delivery^41^. To assess whether these changes in efficiency of Cas9 delivery were the result of differences in cargo loading, EVs were analyzed for Cas9 loading abundance by western blot analysis (Fig. 3I). Indeed, western blot analysis revealed a pattern in Cas9 loading levels that correlated with the activation of eGFP expression levels observed in Cas9 delivery, indicating that the observed differences in Cas9 delivery efficiency of these constructs were mainly due to differences in levels of Cas9 loading. To further characterize the effects of the choice of EV-marker for targeted cargo loading, cells were once more transfected with MCP-PhoCl-CD9, MCP-PhoCl-CD63, and MCP-PhoCl-CD81, alongside VSV-G, MS2-sgRNA and Cas9, but were normalized by the number of EV-producing cells, instead of the number of EVs (Fig. S3A). Similar to previous observations, CD63 and CD81 showed equal levels of Cas9 delivery, and CD9 again showed substantially higher levels of Cas9 delivery. In a further comparison, a dose range from 1.0 x 10^8^ – 2.5 x 10^12^ EVs loaded with Cas9 using either MCP-PhoCl-CD9 or MCP- PhoCl-CD63 was added to ∼10^5^ HEK293T reporter cells. Flow cytometry analysis (Fig. S3B) and fluorescence microscopy (Fig. S3C) both showed dose-dependent activation of the stoplight reporter cells, wherein MCP-PhoCl-CD9 once more substantially outperformed MCP-PhoCl- CD63 at all tested concentrations, reaching up to 57% activation at the highest dose. Taken together, these data show that choice of the EV-enriched marker strongly affects the efficiency of EV-mediated Cas9 delivery across various conditions and dosages.

### Delivery of Cas9 transcriptional activators and base editors

As stated above, we aimed to establish a modular EV loading strategy that can be applied to a variety of Cas9-based effector proteins. To test the versatility of this loading strategy, we tested the loading and delivery of the Cas9-based transcriptional activator dCas9-VPR^10^. Here, the nuclease domains of Cas9 have been deactivated, forming a “dead” Cas9 (dCas9), which no longer induces double stranded DNA breaks. Instead, various transcriptional activators have been fused to this protein, allowing the temporary activation of a gene when targeting its promoter region. To measure dCas9-VPR activity, we generated reporter cell lines that contain a doxycycline inducible eGFP expression construct (Fig. 4A). This construct can be activated by addition of doxycycline, which activates rtTA3 resulting in transcription of the eGFP gene downstream of a Tet Responsive Element (TRE) sequence. Alternatively, an sgRNA targeting the TRE sequence facilitates dCas9-VPR-mediated expression of eGFP. Functionality of this generated reporter construct in HEK293T cells was confirmed by fluorescence microscopy (Fig. 4B) and flow cytometry analysis (Fig. 4C). As activation of eGFP expression is induced in a dose-dependent manner, unlike the previously analyzed stoplight reporter construct, eGFP expression levels are shown in MFI, rather than absolute percentages of cells surpassing a set threshold (Fig. S1B). Indeed, as expected, both analyses confirmed that addition of doxycycline induced high levels of eGFP expression. Moreover, transfection of dCas9-VPR alongside a targeting (T) sgRNA or a T MS2-sgRNA also resulted in a substantial, albeit somewhat lower, level of eGFP expression. As a negative control, no increase in eGFP expression was observed when dCas9-VPR was transfected alongside a non-targeting (NT) sgRNA (Fig. 4C). Next, EVs were isolated from cells transfected with VSV-G, dCas9-VPR, MCP- PhoCl-CD9 or MCP-PhoCl-CD63, and NT MS2-sgRNAs or T MS2-sgRNAs, treated with 395 nm UV for 20 minutes on ice, and added to the HEK293T inducible eGFP reporter lines. 48 hours later, eGFP expression levels were measured by flow cytometry analysis (Fig. 4D), showing a significant increase in MFI (ΔMFI) for both MCP-PhoCl-CD9 and MCP-PhoCl-CD63 EVs when isolated from cells expressing a T MS2-sgRNA, but not when expressing a NT MS2-sgRNA. Once more, EVs loaded with MCP-PhoCl-CD9 showed a substantially higher level of reporter activation than the EVs loaded with MCP-PhoCl-CD63. However, both showed significant levels of increase in eGFP expression as compared to untreated controls. These data confirm that the modular MCP-PhoCl-based loading platform is also suitable for the loading and delivery of the dCas9-VPR transcriptional activator, allowing the temporary transcriptional activation of targeted genes without the introduction of permanent genetic mutations.

**Figure 4.**
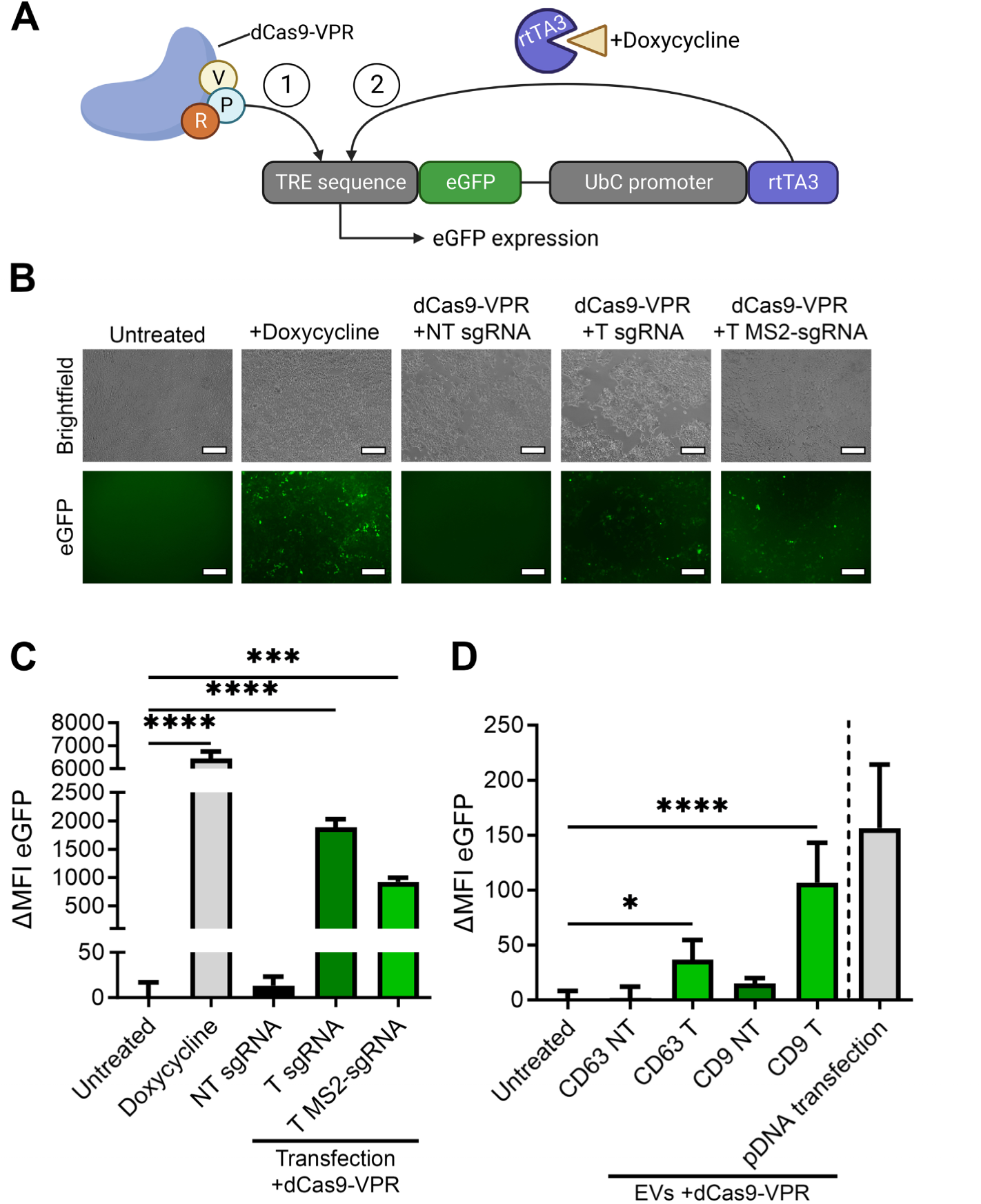
EV-mediated functional delivery of the dCas9-VPR transcriptional activator. **a,** Schematic of the fluorescent reporter construct for transcriptional activation; pInducer20-eGFP. An eGFP open reading frame is placed after a Tet Responsive Element (TRE) sequence. Transcription of eGFP can either be facilitated by activation of the co-expressed reverse tet-transactivator rTA3 by addition of doxycycline (1), or by introduction of transcriptional activator dCas9-VPR with a sgRNA targeting the TRE sequence (2). **b,c** Fluorescence microscopy images **(b)** and flow cytometry analysis **(c)** of HEK293T cells expressing the pInducer20-eGFP reporter construct, 48 hours after addition of doxycycline (0.5 µg/ml), or transfection with plasmids encoding for dCas9-VPR with non-targeting (NT) sgRNAs, targeting (T) sgRNAs, or targeting MS2-sgRNAs. Doxycyline and dCas9-VPR with targeting sgRNAs increase eGFP expression. MFI: mean fluorescence intensity. Scalebar represents 200 μm. Means + SD, n = 3, One-way ANOVA with post-hoc Dunnett’s multiple comparison test. **d,** Flow cytometry analysis of HEK293T cells expressing the pInducer20-eGFP reporter construct, 48 hours after addition of EVs from HEK293T expressing dCas9-VPR alongside either MCP-PhoCl-CD63 or MCP-PhoCl-CD9, in combination with non-targeting- (NT), or targeting (T) sgRNAs. Both loading constructs facilitate EV-mediated functional dCas9- VPR delivery, resulting in a significant increase in eGFP mean fluorescence intensity (MFI). 4.0x10^11^ EVs per well. Means + SD, n = 5, One-way ANOVA with post-hoc Dunnett’s multiple comparison test. * *p* < 0.05, *** *p* < 0.001, **** *p* < 0.0001.

Lastly, we tested the loading and delivery of adenine base editors (ABEs)^12^. Adenine base editors consist of a Cas9 nickase (nCas9), with a DNA modifying deoxyadenosine deaminase (TadA) fused to its N-terminus. This deoxyadenosine deaminase has the capacity to change adenine to inosine, resulting in the conversion of A to G nucleotides within a limited window of the sgRNA targeting sequence. When targeting the DNA strand complementary to the coding strand, this results in the conversion of T to C nucleotides on the coding strand. In order to study ABE delivery, we generated a fluorescent Cas9 adenine base editor (ABE) stoplight reporter construct (Fig. 5A)^17^. In this construct, mCherry is constitutively expressed under a CMV promoter, followed by a stop codon and directly thereafter an in-frame eGFP open reading frame. Through the use of silent mutations, a PAM site was introduced 16 bp upstream of the TAA stopcodon, allowing the targeting and conversion of the stop codon to CAA, encoding for a glutamine, by ABEs. This will result in the expression of an mCherry-eGFP fusion protein, of which the eGFP signal can be measured and quantified by flow cytometry (Fig. S1C). To confirm the functionality of this ABE stoplight reporter, HEK293T reporter cells were generated and transfected with ABE7.10 with non-targeting (NT) or a targeting (T) sgRNA. As a positive control, wildtype Cas9 with a targeting sgRNA with an HDR ssODN template was transfected. As shown by fluorescence microscopy (Fig. S4A) and flow cytometry analysis (Fig. S4B), both ABE7.10 with a targeting sgRNA and WT Cas9 in combination with a T sgRNA and an HDR template were able to activate eGFP expression, albeit at insufficient efficiency to serve as a suitable read-out for EV-mediated ABE delivery (12.9% and 8.5%, respectively). Next, to improve editing efficiency, the more recent ABE8e and ABE8e-dimer proteins were compared to ABE7.10 and analyzed by fluorescence microscopy (Fig. S4C) and flow cytometry analysis (Fig. S4D). Transfection with both ABE8e and ABE8e-dimer resulted in a substantially higher level of eGFP expression, where both induced eGFP expression in over 60% of cells. As the ABE8e and ABE8e-dimer performed similarly, we opted to proceed with ABE8e, due to its smaller size.

**Figure 5.**
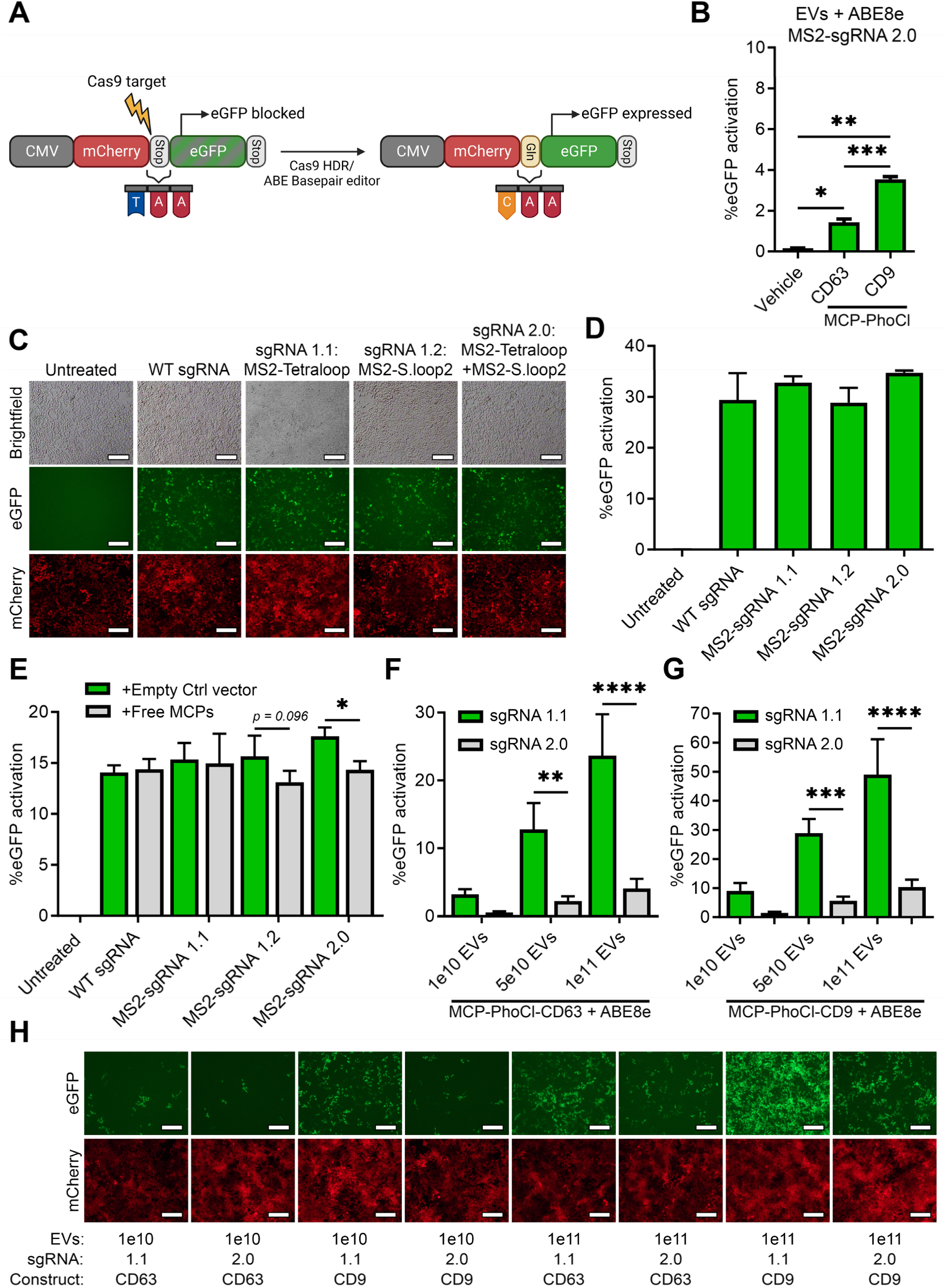
MCP-PhoCl-based EV delivery of adenine base editor ABE8e facilitates high levels of base editing. **a,** Schematic of the fluorescent reporter construct for adenine base editor (ABE) activity. mCherry is expressed under a CMV promotor, followed directly by a stop codon and an in-frame eGFP ORF. Using silent mutations, a Cas9 target site was generated near the mCherry stop codon, allowing ABE-mediated conversion of the stop codon into a glutamine (Gln) codon, resulting in the expression of a mCherry-eGFP fusion protein. **b,** EV- mediated delivery of ABE8e using MCP-PhoCl-CD63 or MCP-PhoCl-CD9 in combination with a targeting MS2- sgRNA 2.0 results in significant but limited ABE8e activity. 5 x 10^10^ EVs per well. Means + SD, n=3, One-way ANOVA with post-hoc Tukey’s multiple comparisons test. **c, d** Transfection of plasmid DNA encoding for ABE8e with a targeting WT sgRNA (no MS2 stemloops), MS2-sgRNA 1.1 (MS2 aptamer in tetraloop), MS2-sgRNA 1.2 (MS2 aptamers in stem loop 2), or MS2-sgRNA 2.0 (MS2 aptamer on tetraloop and stemloop 2) results in similar levels of ABE8e activity, as shown by fluorescence microscopy images **(c)** and flow cytometry analysis **(d)**, 48 hours after transfection. Scalebar represents 200 μm. Means + SD, n = 3. **(e)** Flow cytometry analysis of HEK293T fluorescent reporter cells transfected with ABE8e and various targeting sgRNAs, alongside a plasmid for expression of cytosolic tandem MCPs, or an empty control vector. Only sgRNAs with an MS2 aptamer in the second stemloop show decreased ABE8e activity in the presence of free MCPs. Means + SD, n = 3, One-way ANOVA with post-hoc Sidak’s multiple comparison test **f, g, h** Flow cytometry analysis **(f, g)** and fluorescence microscopy images **(h)** for a dose response of EV-mediated ABE8e delivery using MCP-PhoCl-CD63 **(f)** and MCP- PhoCl-CD9 **(g)** comparing MS2-sgRNA 1.1 and 2.0. Both loading constructs show high ABE8e activity with MS2- sgRNA 1.1, in a dose-dependent manner. Scalebar represents 200 μm. Means + SD, n = 3, One-way ANOVA with post-hoc Sidak’s multiple comparison test. * *p* < 0.05, ** *p* < 0.01, *** *p* < 0.001, **** *p* < 0.0001.

Next, EVs were isolated from HEK293T cells expressing VSV-G, MS2-sgRNA, ABE8e, and MCP- PhoCl-CD63 or MCP-PhoCl-CD9, were treated with 395 nm UV and added to our ABE stoplight reporter cells. Disappointingly, both constructs showed fairly low percentages of eGFP activation, around 1.4% and 3.5% respectively. Since the deoxyadenosine deaminase fused to nCas9 has to be able to interact with the DNA to induce the A to G mutations, and the MCPs are still fused to the MS2 aptamers on the MS2-sgRNAs after UV-mediated cleavage from the tetraspanins, we hypothesized that the MCP domains might physically block proper access for the deoxyadenosine deaminase to the DNA. Thus, we generated targeting MS2-sgRNAs with MS2 aptamers only in the Tetraloop (MS2-sgRNA 1.1), only in the second stemloop, (MS2- sgRNA 1.2), in both (MS2-sgRNA 2.0), or in neither (WT sgRNA) (terminology in line with the manuscript from Konermann et al.^33^). To assess the effect of the MS2 aptamers on ABE8e functionality, plasmids for each of these sgRNAs were co-transfected with a plasmid expressing ABE8e into ABE stoplight reporter cells. Flow cytometry analysis showed no significant difference in eGFP expression between these sgRNAs, indicating that the MS2 aptamers do not interfere with ABE8e functionality (Fig. 5D). However, when an additional plasmid expressing free cytosolic tandem MCPs was co-transfected, a decrease in functionality was observed for MS2-sgRNA 1.2 and MS2-sgRNA 2.0, indicating that binding of tandem MCPs to the second stemloop of the sgRNA interferes with ABE8e functionality. To test whether this indeed negatively affected our EV-mediated delivery of ABE8e using MCP- PhoCl constructs, EVs were isolated from HEK293T cells expressing VSV-G, ABE8e, MS2-sgRNA 1.1 or MS2-sgRNA 2.0, in combination with MCP-PhoCl-CD63 or MCP-PhoCl-CD9. EVs were treated with 395 nm UV, and added to 10^5^ ABE Stoplight reporter cells at multiple dosages (1.0 x 10^10^, 5.0 x 10^10^, 1.0 x 10^11^). As previously observed (Fig. 5B), addition of EVs with ABE8e loaded with MS2-sgRNAs with aptamers in both the tetraloop and second stemloop (MS2- sgRNA 2.0) resulted in low levels of eGFP as shown by flow cytometry analysis for MCP-PhoCl- CD63 (Fig. 5F) and MCP-PhoCl-CD9 (Fig. 5G), as well as by fluorescence microscopy (Fig. 5H). However, EV-mediated delivery of ABE8e using MS2-sgRNA 1.1 (tetraloop MS2 aptamer only), resulted in high levels of dose-dependent ABE8e delivery, confirming that MCP binding to the second stemloop of the sgRNA protruding from the RNP indeed inhibited ABE activity. Once more, MCP-PhoCl-CD9 outperformed MCP-PhoCl-CD63 >2-fold, in line with observations for WT Cas9 (Fig. 3G, S3A, S3B) and dCas9-VPR (Fig. 4D). These data show that this platform is also suitable for the delivery of adenine base editors, but MS2 aptamer-MCP binding on the second stemloop of the sgRNA has a strong negative effect on ABE8e functionality, and should thus be avoided.

Altogether, these data show that MCP-PhoCl-mediated EV loading and release constructs provide a modular, versatile EV-mediated Cas9 delivery platform that allows for the efficient delivery of various Cas9-based proteins with high therapeutic potential, including wildtype Cas9, transcriptional activators, and adenine base editors.

## Discussion

Already in the first report where the CRISPR-Cas9 system was harnassed for targeted genomic engineering of specific sequences in prokaryotic cells by Jinek et al.^2^, the authors envisioned the potential of a “methodology based on RNA-programmed Cas9 that could offer considerable potential for gene-targeting and genome-editing applications”. This study also included the first report of a chimeric RNA, an RNA molecule generated by fusing the 3’ end of crRNA to the 5’ end of tracrRNA, laying the foundation for the widely adapted use of single guide RNAs (sgRNAs). Soon after, a study followed that reported the inclusion of a nuclear localization signal to ensure nuclear compartmentalization, making Cas9 suitable for efficient engineering in a wide range of mammalian cells^3^. Hereafter, various publications followed that provided accesible cloning strategies with high ease-of-use, substantially lowering the barrier of entry for the application of CRISPR-Cas9 technology in biomedical research^42,43^. Since these impactful developments, substantial effort has been invested in the design of Cas9-based therapeutic strategies for the treatment of genetic diseases. Promising current clinical trials for direct in vivo Cas9 gene editing strategies are based on lipid nanoparticle (LNP)-mediated Cas9 delivery for the treatment of hereditary angioedema and for transthyretin amyloidosis, by disrupting liver expression of KLKB1 or TTR, respectively^44,45^. Whereas these LNP-based strategies have shown clinical promise for in vivo delivery of Cas9, their restricted biodistribution profile currently limits their application to gene editing strategies specifically targeting the liver. Moreover, LNPs have been reported to show cellular toxicity and limited delivery efficiency due to endosomal entrapment^20^. Furthermore, recent studies have shown that repeated injections with LNPs can lead to antibody-mediated immune reactions against the polyethylene glycol (PEG) groups present on the exterior of LNPs, which may result in accelerated blood clearance and decrease in effectivity^46^. Thus, there is a pressing need for the development of novel delivery strategies with tunable biodistribution profiles, low toxicity, and low immunogenicity for the in vivo delivery of CRISPR-Cas9.

In this study, we describe a modular strategy for extracellular vesicle (EV)-mediated delivery of CRISPR-Cas9, by employing an aptamer-based strategy for loading of Cas9 RNPs into EVs. This approach is combined with the incorporation of a photocleavable domain (PhoCl) to facilitate intraluminal cargo release from the EV membrane after stimulation with 400 nm UV light. We chose to study the suitability of EVs for intracellular Cas9 RNP delivery due to their natural capacity to transfer proteins, RNA, and various other biological cargos as part of their role in intercellular communication^47^. Alongside their intrinsic capacity to transfer biological cargos, EVs are promising delivery vectors due to their reported low immunogenicity, and their capacity to pass hard-to-cross biological barriers such as the blood-brain-barrier^48,49^. To facilitate active Cas9 RNP loading into EVs, we made use of the capacity of MS2 coat proteins (MCPs) to strongly bind to MS2 RNA aptamers^31^. In this system, tandem MCPs bind a single aptamer. By intraluminal fusion of MCPs to proteins that are enriched in EVs, RNA molecules are actively loaded into EVs during their biogenesis. This strategy has previously been reported to facilitate mRNA loading into EVs in an approach called Targeted and Modular EV Loading (TAMEL) by Hung et al.^50^. Here, a 40-fold increase in mRNA loading into EVs was observed when directly fusing single MCP domains directly to VSV-G. Whereas substantial loading was observed in this study, no translation of mRNA cargo was observed in recipient cells. It should be noted that this MCP used contained the V29I mutation, which has been reported to substantially increase MCP-MS2 aptamer affinity^51^. As such, a limiting factor in their delivery strategy may have been the retention of their mRNA cargo to the EV membrane, preventing release into the cytosol of the recipient cell. Here, we applied this strategy the load Cas9 RNPs, based on the incorporation of MS2 aptamers into the tetraloop and second stemloop of the sgRNA^33^. Despite the exclusion of the V29I mutation in the MCP, we still saw limited functional RNP delivery initially. Only after including an activatable release strategy by incorporating the PhoCl photocleavable domain^32^ did we observe a substantial increase in RNP delivery, confirming that cargo retention to the EV membrane was indeed hampering functional delivery.

The PhoCl protein applied in this manuscript is a derivative from the mMaple fluorescent protein, a green fluorescent protein that undergoes a conformational change after UV stimulation, resulting in a switch to red fluorescence^52^. The PhoCl platform is based on the incorporation of various mutations, uncovered by Zhang et al., resulting in protein instability following this UV-mediated transformational change, resulting in protein cleavage^32^. In recent years, various additional strategies have been described to release cargos from the EV membrane to increase cargo delivery. Another light-based cargo loading and release strategy, is the “exosomes for protein loading via optically reversible protein–protein interactions”(EXPLORs) platform^53^. This system is based on the incorporation of photoreceptor cryptochrome 2 (CRY2) on the N-terminus of CD9, and a truncated CRY-interacting basic-helix-loop-helix 1 protein module (CIBN) on the cargo protein. Upon stimulation with blue light, CRY2 binds to the CIBN module, resulting in EV cargo loading. Upon removal of the blue light source, this interaction is abbrogated, resulting in cargo release. Whereas this approach has shown promising efficiency in cargo delivery, one limitation of this system is the requirement of constant blue light exposure throughout cell culture conditions, prior to EV isolation. Another reported strategy is the FK506 binding protein (FKBP) and the FKBP-rapamycin-binding domain (FRB) dimerization system^54,55^. Both these moieties bind to rapamycin analogs, and by fusing FKBP to an EV-enriched protein, and FRB to the cargo protein, cargo loading can be facilitated by addition of rapamycin analogs to cell culture conditions. As these membrane-permeable rapamycin analogs are removed from the surrounding media throughout EV isolation protocols, cargo may be untethered from the EV membrane. One limitation of this system is the requirement of the addition of rapamycin- analogues to cell culture conditions to facilitate cargo loading, resulting in increased costs of EV production. For further examples, we recommend a recent study from Osteikoetxea et al., which shows a thorough comparison of these systems, alongside various additional loading and release strategies for EV-mediated Cas9 delivery^56^. As compared to these stragegies, an advantage of the PhoCl-based release strategy presented in this manuscript is the lack of any additional conditional requirements during cell culture and EV production, as the photocleavable domain is intact until stimulated with UV light. One recently published strategy that has a similar advantage, is the incorporation of self-cleaving inteins^57^. These protein domains can either self-excise themselves from protein sequences or, after specific amino acid substitutions facilitate protein cleavage, may be activated by a variety of conditions including changes in pH, temperature, or naturally over time^58^. One key challenge with intein-based strategies is to incorporate one with optimal kinetics, which may differ per application or even loading pathway. If cleavage occurs too rapidly, cargo loading may be compromised. Conversely, if cleavage occurs too slowly cargo may not be fully released in recipient cells. As such, incorporating pH-dependent inteins as described by Liang et al. is an attractive strategy, as it facilitates cargo release in late endosomal compartments in recipient cells^57^.

Our PhoCl-mediated approach also faces certain limitations, as UV-mediated cleavage occurs at a fairly long half-time (∼500 seconds) and dissociation efficiency is limited to ∼71%^32^. However, recent work from Lu et al. has demonstrated novel PhoCl derivatives that either show increased cleavage rates (PhoCl2f, 76 second half-time) and increased efficiency (PhoCl2c, 92% dissociation efficiency)^59^, which may further improve the efficiency and ease-of-use of the delivery platform described in this study. Another point of initial concern was the potential damage to EV components or its Cas9 RNP cargo resulting from UV treatment. However, it should be noted that UV wavelengths associated with nucleotide damage are below 340 nm^60^. Concordantly, we also did not observe any decrease in Cas9 RNP delivery after applying 395 nm UV treatment to EVs that did not contain the PhoCl domain, indicating that the Cas9 RNP cargo indeed remained intact. However, we cannot completely exclude potential effects of our UV treatment on EV integrity or functionality. As such, the use of the aforementioned PhoCl2f photocleavable domain with highly increased cleavage rates might be of interest to further reduce this risk in future studies.

For active cargo loading, our current approach relied on the MCP-MS2 aptamer interaction. Whereas this system is highly efficient and well-designed it did present some challenges, most notably in terms of cargo release. Whereas this was, in part, alleviated by the incorporation of a photocleavable domain, the tandem MCP complex is still bound to the sgRNA MS2 aptamers on the RNP complex. In line with previous reports, this did not compromise the functionality of Cas9^33^. However, we did observe interference with the functionality of adenine base editor ABE8e. As this was only observed when incorporating MS2 aptamers into the second stemloop of the sgRNA we hypothesize that the reduction in functionality is caused by interfering with the interaction of the TadA domain with the targeted DNA sequence, as the second stemloop is relatively close to the TadA domain^61^. Whereas this issue can be addressed by only incorporating MS2 aptamers into the tetraloop of the sgRNA, it does reveal a limitation in the robustness of the modular versatility of this approach. A potential solution may be found in the utilisation of different aptamer-based loading systems that rely on smaller RNA-binding proteins such as the COM aptamers system, or that show differences in release kinetics such as the recently reported optimized version of the designer Pumilio and FBF homology domain, termed PUFe^62,63^. Lastly, we also observed substantial differences in Cas9 RNP delivery based on the choice of EV-enriched protein used for targeted loading. Interestingly, whereas CD9 significantly outperformed CD63 in terms of both cargo loading and delivery in our study, a recent study from Zheng et al. showed substantially higher loading when using CD63 as compared to CD9^29^. One potential explanation for this observed difference might be that whereas our cargo loading strategy was based on N-terminal fusion to the EV-enriched proteins, Zheng et al. opted for C-terminal fusion given that N-termini are often a site for signal peptides that affect intracellular trafficking. As, to the best of our knowledge, no direct comparison has been done on the effects of C-terminal and N-terminal fusion on tetraspanins for EV-loading efficiency as of yet, taking this additional parameter into consideration may be warranted for future design strategies. Furthermore, efficiency of our delivery strategy may be further improved by incorporating recently uncovered moieties that are highly enriched in EVs, including TSPAN2, PTGFRN, a MysPalm tag, or direct fusion to the VSV-G glycoprotein^29,30,50,56^.

In closing, we describe a modular, versatile and efficient engineering strategy for EV-based delivery of Cas9, based on the combined utilization of aptamer-based RNP loading and UV-activated cargo release. This strategy is suitable for the delivery of various Cas9 variants, facilitating NHEJ-based gene editing, transcriptional activation, and adenine base editing. Given the modular design of this approach, there is strong potential for additional future applications, such as the delivery of mRNA, CRISPR-mediated transcriptional inhibitors (CRISPRi), and additional base editing approaches such as prime- and cytidine base editing. Altogether, this work further broadens the utility of EVs for (bio)therapeutic delivery strategies, and underlines the potential for EV-mediated treatment of genetic diseases.

## Acknowledgements

O.G.d.J. was supported by a VENI Fellowship (VI.Veni.192.174) from the Dutch Research Council (NWO). The work of O.E, R.M.S. and P.V. is supported by the European Union’s Horizon 2020 Research and Innovation Programme under grant agreement No. 825828 (EXPERT). W.S.d.V. and P.V. are supported by the European Research Council (ERC) Starting grant OBSERVE (No. 851936).C.V.H. was supported by the Utrecht Institute for Pharmaceutical Sciencess (UIPS). Figures 1A, 2A, 3A, 3F, 4A and 5A were generated using Biorender.com.

## Author contributions

O.G.d.J and P.V. initiated this project. S.E.A., R.M.S., E.M., and S.A.A. helped design and plan the project. O.G.d.J., P.V., and O.E. planned and designed the experiments. C.V.H., I.L., O.L.C, I.Y.d.G., Z.E.N.M.J.d.W., X.L., A.G.G., N.J.A.M., J.L., W.S.d.V., J.J.G.F., and A.C.W.v.W. helped design and perform the experiments. O.G.d.J. and O.E. designed and developed most of the methodology. O.G.d.J. was responsible for the overall project strategy and management and wrote the manuscript, which was reviewed by all authors.

