## Supplementary Data for "A modular strategy for extracellular vesicle-mediated CRISPR-Cas9 delivery through aptamer-based loading and UV-activated cargo release"

### Supplementary Figures

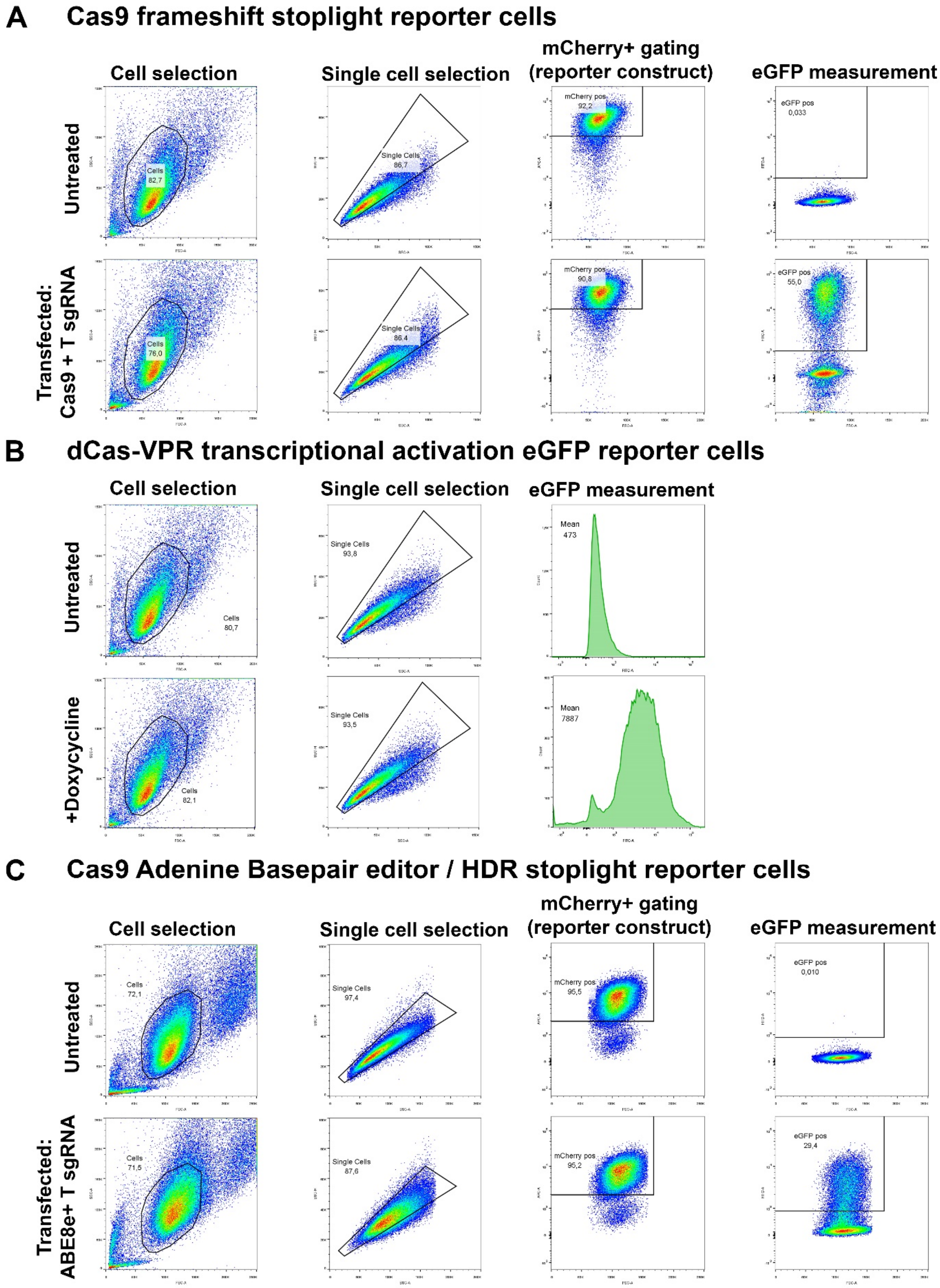

**Supplementary Figure 1. Flow cytometry gating strategy employed for the analysis of fluorescent Cas9 activity reporter constructs. a,** Gating strategy for analysis of the “stoplight” reporter construct for Cas9 activity-mediated frameshifts (schematic: Fig. 2A). Cells were gated using the forward scatter (FSC-A) and sideward scatter (SSC-A) signals. Single cells were selected using sideward scatter area (SSC-A) and sideward scatter height (SSC-H) signals. Next, mCherry+ cells were selected using FSC-A and mCherry signals to ensure stoplight reporter expression. Lastly, eGFP expression was measured within mCherry+ cells using FSC-A and eGFP signals. Representative plots are shown for untreated HEK293T reporter cells (top row), and HEK293T reporter cells transfected with plasmid DNA for expression of Cas9 and a targeting sgRNA (bottom row). **b,** Gating strategy for analysis of the transcriptional activator reporter construct, based on inducible eGFP expression; pInducer20, dCas9-VPR activity (schematic: Fig. 4A). Cells were gated using the forward scatter (FSC-A) and sideward scatter (SSC-A) signals. Single cells were selected using sideward scatter area (SSC-A) and sideward scatter height (SSC-H) signals. eGFP signal was then measured and mean fluorescence intensity (MFI) was determined. Representative plots are shown for untreated HEK293T reporter cells (top row), and HEK293T reporter cells treated with doxycycline (positive control, bottom row). **c,** Gating strategy for analysis of the “stoplight” reporter construct for adenine base editor (ABE) activity (schematic: Fig. 5A). Cells were gated using the forward scatter (FSC-A) and sideward scatter (SSC-A) signals. Single cells were selected using sideward scatter area (SSC-A) and sideward scatter height (SSC-H) signals. Next, mCherry+ cells were selected using FSC-A and mCherry signals to ensure stoplight reporter expression. Lastly, eGFP expression was measured within mCherry+ cells using FSC-A and eGFP signals. Representative plots are shown for untreated HEK293T reporter cells (top row), and HEK293T reporter cells transfected with plasmid DNA for expression of ABE8e and a T sgRNA (bottom row).

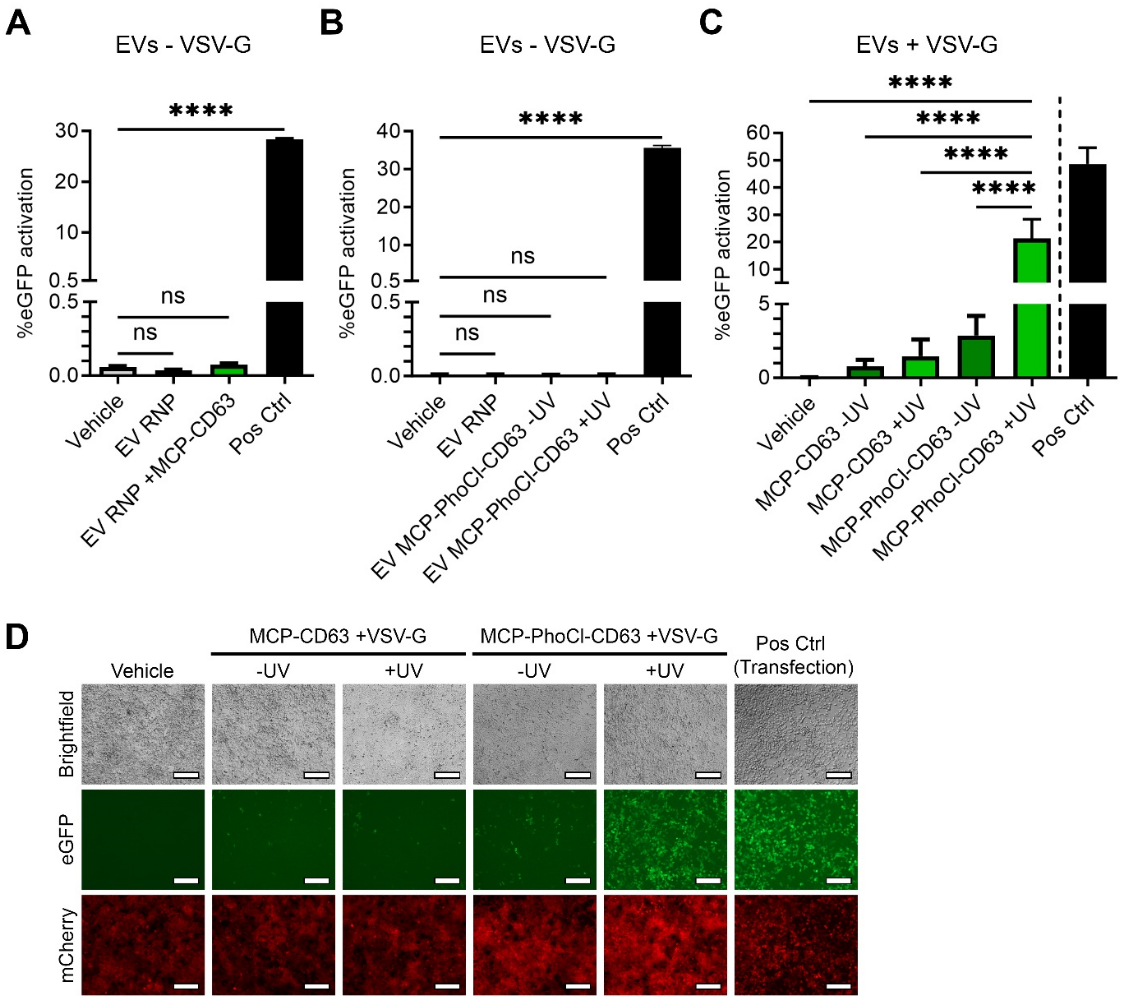


**Supplementary Figure 2. Effects of VSV-G and UV on MCP-CD63 and MCP-PhoCl-CD63 Cas9 delivery. a,** Flow cytometry analysis of HEK293T cells expressing the stoplight reporter construct for Cas9 activity 72 hours after addition of EVs isolated from HEK293T cells expressing Cas9 + MS2-sgRNA (EV RNP) or expressing Cas9 + MS2-sgRNA + MCP-CD63 (EV RNP + MCP-CD63) without the co-expression of VSV-G shows no detectable EV-mediated functional Cas9 delivery. Means + SD, n = 3, One-way ANOVA with Dunnett's multiple comparison test. **b,** Flow cytometry analysis of HEK293T cells expressing the stoplight reporter construct for Cas9 activity 72 hours after addition of EVs isolated from HEK293T cells expressing Cas9 + MS2-sgRNA (EV RNP) or expressing Cas9 + MS2-sgRNA + MCP-PhoCl-CD63 with- (+UV) or without (-UV) UV treatment, without the co-expression of VSV-G shows no detectable EV-mediated functional Cas9 delivery. **c, d** Flow cytometry analysis **(c)** and fluorescence microscopy images **(d)** of HEK293T cells expressing the stoplight reporter construct for Cas9 activity 72 hours after addition of EVs isolated HEK293T cells expressing VSV-G, Cas9 and a targeting MS2-sgRNA alongside MCP-CD63 or MCP-PhoCl-CD63, either treated with- or without UV. MCP-PhoCl-CD63 with UV significantly outperforms all other conditions. UV treatment does not result in decreased Cas9 delivery in MCP-CD63 EVs, indicating that cargo is not negatively affected by UV. Means + SD, n = 3, One-way ANOVA with Tukey's multiple comparison test. Scalebar represents 200 µm. **** $p < 0.0001$.

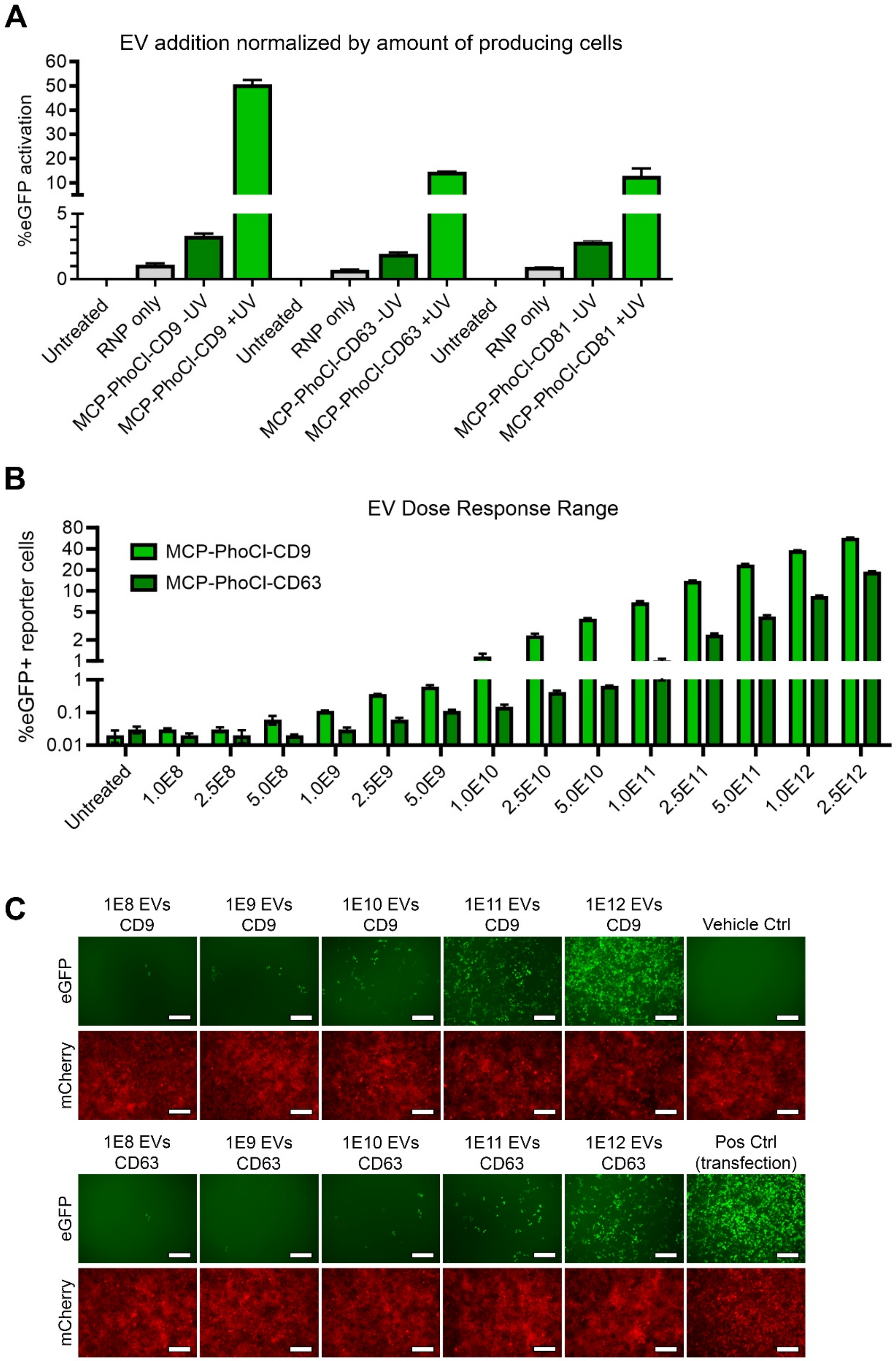


**Supplementary Figure 3. Comparison of tetraspanins in MCP-PhoCl constructs normalized on cell count and particle count. a,** Comparison of MCP-CD9-PhoCl, MCP-CD63-PhoCl, and MCP-CD81-PhoCl normalized on cell count of EV-producing cells co-expressing VSV-G, Cas9 and MS2-sgRNA. EVs from $5x10^7$ cells were added per well. Relative differences in Cas9 delivery efficiency between CD9, CD63 and CD81 is comparable to experiments normalized to particle count (shown in Fig. 3). Means + SD, n = 3. **b, c** Flow cytometry analysis **(b)**

and fluorescence microscopy images **(c)** of HEK293T cells expressing the stoplight reporter construct for Cas9 activity 72 hours after addition of an EV dose-range, between 1.0x$10^8$ and 2.5x$10^{12}$ EVs per well, from HEK293T cells expressing VSV-G, Cas9 and a targeting MS2-sgRNA alongside MCP-PhoCl-CD9 and MCP-PhoCl-CD63, confirming a dose-dependent delivery of Cas9. Scale bar represents 200 µm. Means + SD, n = 3.

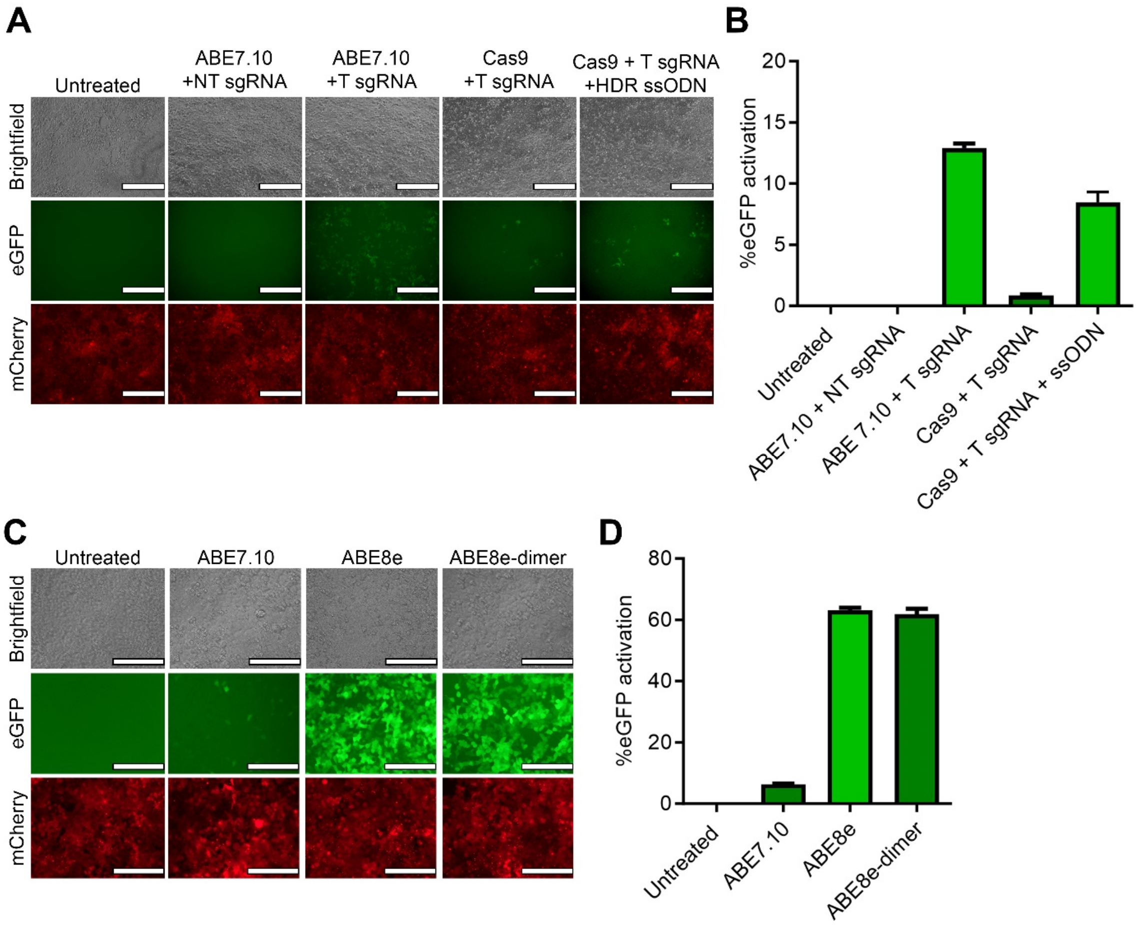


**Supplementary Figure 4. Cas9-mediated homology directed repair and adenine base editor comparison on fluorescent adenine base editor reporter. a,b** Fluorescent microscopy images **(a)** and flow cytometry analysis **(b)** of HEK293T cells expressing the stoplight reporter construct for ABE activity, 72 hours after transfection with plasmid DNA expressing ABE7.10 with a non-targeting (NT) or targeting (T) sgRNA, or Cas9 with a T sgRNA with- or without a ssODN HDR template. Transfection of both ABE7.10 and Cas9 with an ssODN template with their T sgRNAs induce eGFP expression. Means + SD, n = 3. **c, d** Fluorescent microscopy images **(c)** and flow cytometry analysis **(d)** of HEK293T cells expressing the stoplight reporter construct for ABE activity, 96 hours after transfection with plasmid DNA expressing ABE7.10, ABE8e or ABE8e-dimer with a T sgRNA. ABE8e and ABE8e-dimer show similar activity, and both strongly outperform ABE7.10. Scale bar represents 200 µm. Means + SD, n = 3.

### Supplementary tables

##### Supplementary Table 1: Ordered oligonucleotide sequences

| Target | Purpose | Orientation | Sequence |
|---|---|---|---|
| Stoplight | sgRNA cloning | Sense | 5’-CACCGGGACAGTACTCCGCTCGAGT-3’ |
| Stoplight | sgRNA cloning | Antisense | 5’-AAACACTCGAGCGGAGTACTGTCCC-3 |
| Tet resp. element | sgRNA cloning | Sense | 5’-CACCGGTCTCTATCACTGATAGGGAG-3’ |
| Tet resp. element | sgRNA cloning | Antisense | 5’-AAACCTCCCTATCAGTGATAGAGACC-3’ |
| ABE Stoplight | sgRNA cloning | Sense | 5’-CACCGCTTACTTGTACAGCTCGTCC-3’ |
| ABE Stoplight | sgRNA cloning | Antisense | 5’-AAACGGACGAGCTGTACAAGTAAGC-3’ |
| ABE Stoplight | HDR template | Antisense | 5’-ACGACGCCCGTGAAAAGCTCTTCACCCTTAGACACGGCTTGCTTGTACAGCTCGTCCAAGCCGCCCGTAGAATGCCTGCCT-’3 |

##### Supplementary Table 2: Expressed sgRNA sequences

| Target | sgRNA | Sequence |
|---|---|---|
| Stoplight | WT | GGACAGUACUCCGCUCGAGUGUUUUAGAGCUAGAAAUAGCAAGUUAAAAUAAGGCUAGUCCGUUAUCAACUUGAAAAAGUGGCACCGAGUCGGUGCUUUUUU |
| Stoplight | MS2-2.0 | GGACAGUACUCCGCUCGAGUGUUUUAGAGCUAGGCCAACAUGAGGAUCACCCAUGUCUGCAGGGCCUAGCAAGUUAAAAUAAGGCUAGUCCGUUAUCAACUUGGCCAACAUGAGGAUCACCCAUGUCUGCAGGGCCAAGUGGCACCGAGUCGGUGCUUUUUUU |
| Tet resp. element | WT | GUCUCUAUCACUGAUAGGGAGGUUUUAGAGCUAGAAAUAGCAAGUUAAAAUAAGGCUAGUCCGUUAUCAACUUGAAAAAGUGGCACCGAGUCGGUGCUUUUUU |
| Tet resp. element | MS2-2.0 | GUCUCUAUCACUGAUAGGGAGGUUUUAGAGCUAGGCCAACAUGAGGAUCACCCAUGUCUGCAGGGCCUAGCAAGUUAAAAUAAGGCUAGUCCGUUAUCAACUUGGCCAACAUGAGGAUCACCCAUGUCUGCAGGGCCAAGUGGCACCGAGUCGGUGCUUUUUUU |
| ABE Stoplight | WT | GCUUACUUGUACAGCUCGUCCGUUUUAGAGCUAGAAAUAGCAAGUUAAAAUAAGGCUAGUCCGUUAUCAACUUGAAAAAGUGGCACCGAGUCGGUGCUUUUUU |
| ABE Stoplight | MS2-1.1 | GCUUACUUGUACAGCUCGUCCGUUUUAGAGCUAACAUGAGGAUCACCCAUGUUAGCAAGUUAAAAUAAGGCUAGUCCGUUAUCAACUUGGACUUCGGUCCAAGUGGCACCGAGUCGGUGCUUUUUUU |
| ABE Stoplight | MS2-1.2 | GCUUACUUGUACAGCUCGUCCGUUUCAGAGCUAAGCACAAGAGUGCAUAGCAAGUUGAAAUAAGGCUAGUCCGUUUACAACUUGGCCAACAUGAGGAUUACCCAUGUCUGCAGGGCCAAGUGGCACCCGAGUCGGGUGCUUUUUUU |
| ABE Stoplight | MS2-2.0 | GCUUACUUGUACAGCUCGUCCGUUUUAGAGCUAGGCCAACAUGAGGAUCACCCAUGUCUGCAGGGCCUAGCAAGUUAAAAUAAGGCUAGUCCGUUAUCAACUUGGCCAACAUGAGGAUCACCCAUGUCUGCAGGGCCAAGUGGCACCGAGUCGGUGCUUUUUUU |

Legend: sgRNA targeting sequence, MS2 aptamer.

##### Supplementary Table 3: Amino acid sequences of RNA-binding constructs

| Construct | Amino acid sequence |
|---|---|
| MCP-CD63 | MASNFTQFVLVDNGGTGDVTVAPSNFANGVAEWISSNSRSQAYKVTCSVRQSSAQNRKYTIKVEVPKGAWRSYLNMELTIPIFATNSDCELIVKAMQGLLKDGNPIPSAIAANSGIYAMASNFTQFVLVDNGGTGDVTVAPSNFANGVAEWISSNSRSQAYKVTCSVRQSSAQNRKYTIKVEVPKGAWRSYLNMELTIPIFATNSDCELIVKAMQGLLKDGNPIPSAIAANSGIYGSSPSTSLYKKAGSEFALKLAVEGGMKCVKFLLYVLLLAFCACAVGLIAVGVGAQLVLSQTIIQGATPGSLLPVVIIAVGVFLFLVAFVGCCGACKENYCLMITFAIFLSLIMLVEVAAAIAGYVFRDKVMSEFNNNFRQQMENYPKNNHTASILDRMQADFKCCGAANYTDWEKIPSMSKNRVPDSCCINVTVGCGINFNEKAIHKEGCVEKIGGWLRKNVLVVAAAALGIAFVEVLGIVFACCLVKSIRSGYEVM* |

| | |
|---|---|
| MCP-PhoCl-CD63 | MASNFTQFVLVDNGGTGDVTVAPSNFANGVAEWISSNSRSQAYKVTCSVRQSSAQNRKYT<br>IKVEVPKGAWRSYLNMELTIPIFATNSDCELIVKAMQGLLKDGNPIPSAIAANSGIYAMASNF<br>TQFVLVDNGGTGDVTVAPSNFANGVAEWISSNSRSQAYKVTCSVRQSSAQNRKYTIKVEVP<br>KGAWRSYLNMELTIPIFATNSDCELIVKAMQGLLKDGNPIPSAIAANSGIYGSSYPYDVPDYA<br>VIPDYFKQSFPEGYSWERSMTYEDGGICIATNDITMEGDSFINKIHFKGTNFPPNGPVMQKR<br>TVGWEASTEKMYERDGVLKGDVKMKLLLKGGGHYRCDYRTTYKVKQKPVKLPDYHFVDHRI<br>EILSHDKDYNKVKLYEHAVARNSTDSMDELYKGGSGGMVSKGEETITSVIKPDMKNKLRME<br>GNVNGHAFVIEGEGSGKPFEGIQTIDLEVKEGAPLPFAYDILTTAFHYGNRVFTKYPRDYKDD<br>DDKLYKKAGSEFALKLAVEGGMKCVKFLLYVLLLAFCACAVGLIAVGVGAQLVLSQTIIQGAT<br>PGSLLPVVIIAVGVFLFLVAFVGCCGACKENYCLMITFAIFLSLIMLVEVAAAIAGYVFRDKVMS<br>EFNNNFRQQMENYPKNNHTASILDRMQADFKCCGAANYTDWEKIPSMSKNRVPDSCCIN<br>VTVGCGINFNEKAIHKEGCVEKIGGWLRKNVLVVAAAALGIAFVEVLGIVFACCLVKSIRSGYE<br>VM* |
| MCP-PhoCl-CD9 | MASNFTQFVLVDNGGTGDVTVAPSNFANGVAEWISSNSRSQAYKVTCSVRQSSAQNRKYT<br>IKVEVPKGAWRSYLNMELTIPIFATNSDCELIVKAMQGLLKDGNPIPSAIAANSGIYAMASNF<br>TQFVLVDNGGTGDVTVAPSNFANGVAEWISSNSRSQAYKVTCSVRQSSAQNRKYTIKVEVP<br>KGAWRSYLNMELTIPIFATNSDCELIVKAMQGLLKDGNPIPSAIAANSGIYGSSYPYDVPDYA<br>VIPDYFKQSFPEGYSWERSMTYEDGGICIATNDITMEGDSFINKIHFKGTNFPPNGPVMQKR<br>TVGWEASTEKMYERDGVLKGDVKMKLLLKGGGHYRCDYRTTYKVKQKPVKLPDYHFVDHRI<br>EILSHDKDYNKVKLYEHAVARNSTDSMDELYKGGSGGMVSKGEETITSVIKPDMKNKLRME<br>GNVNGHAFVIEGEGSGKPFEGIQTIDLEVKEGAPLPFAYDILTTAFHYGNRVFTKYPRDYKDD<br>DDKLYKKAGSEFALKLPVKGGTKCIKYLLFGFNFIFWLAGIAVLAIGLWLRFDSQTKSIFEQETN<br>NNNSSFYTGVYILIGAGALMMLVGFLGCCGAVQESQCMLGLFFGFLLVIFAIEIAAAIWGYSH<br>KDEVIKEVQEFYKDTYNKLKTKDEPQRETLKAIHYALNCCGLAGGVEQFISDICPKKDVLETFT<br>VKSCPDAIKEVFDNKFHIIGAVGIGIAVVMIFGMIFSMILCCAIRRNREMV* |
| MCP-PhoCl-CD81 | MASNFTQFVLVDNGGTGDVTVAPSNFANGVAEWISSNSRSQAYKVTCSVRQSSAQNRKYT<br>IKVEVPKGAWRSYLNMELTIPIFATNSDCELIVKAMQGLLKDGNPIPSAIAANSGIYAMASNF<br>TQFVLVDNGGTGDVTVAPSNFANGVAEWISSNSRSQAYKVTCSVRQSSAQNRKYTIKVEVP<br>KGAWRSYLNMELTIPIFATNSDCELIVKAMQGLLKDGNPIPSAIAANSGIYGSSYPYDVPDYA<br>VIPDYFKQSFPEGYSWERSMTYEDGGICIATNDITMEGDSFINKIHFKGTNFPPNGPVMQKR<br>TVGWEASTEKMYERDGVLKGDVKMKLLLKGGGHYRCDYRTTYKVKQKPVKLPDYHFVDHRI<br>EILSHDKDYNKVKLYEHAVARNSTDSMDELYKGGSGGMVSKGEETITSVIKPDMKNKLRME<br>GNVNGHAFVIEGEGSGKPFEGIQTIDLEVKEGAPLPFAYDILTTAFHYGNRVFTKYPRDYKDD<br>DDKLYKKAGSEFALKLGVEGCTKCIKYLLFVFNFVFWLAGGVILGVALWLRHDPQTTNLLYLEL<br>GDKPAPNTFYVGIYILIAVGAVMMFVGFLGCYGAIQESQCLLGTFFTCLVILFACEVAAGIWG<br>FVNKDQIAKDVKQFYDQALQQAVVDDDANNAKAVVKTFHETLDCCGSSTLTALTTSVLKNN<br>LCPSGSNIISNLFKEDCHQKIDDLFSGKLYLIGIAAIVVAVIMIFEMILSMVLCCGIRNSSVY* |
| MCP-PhoCl-ARRDC1 | MASNFTQFVLVDNGGTGDVTVAPSNFANGVAEWISSNSRSQAYKVTCSVRQSSAQNRKYT<br>IKVEVPKGAWRSYLNMELTIPIFATNSDCELIVKAMQGLLKDGNPIPSAIAANSGIYAMASNF<br>TQFVLVDNGGTGDVTVAPSNFANGVAEWISSNSRSQAYKVTCSVRQSSAQNRKYTIKVEVP<br>KGAWRSYLNMELTIPIFATNSDCELIVKAMQGLLKDGNPIPSAIAANSGIYGSSYPYDVPDYA<br>VIPDYFKQSFPEGYSWERSMTYEDGGICIATNDITMEGDSFINKIHFKGTNFPPNGPVMQKR<br>TVGWEASTEKMYERDGVLKGDVKMKLLLKGGGHYRCDYRTTYKVKQKPVKLPDYHFVDHRI<br>EILSHDKDYNKVKLYEHAVARNSTDSMDELYKGGSGGMVSKGEETITSVIKPDMKNKLRME<br>GNVNGHAFVIEGEGSGKPFEGIQTIDLEVKEGAPLPFAYDILTTAFHYGNRVFTKYPRDYKDD<br>DDKLYKKAGSEFALKLMGRVQLFEISLSHGRVVYSPGEPLAGTVRVRLGAPLPFRAIRVTCIGS<br>CGVSNKANDTAWVVEEGYFNSSLSLADKGSLPAGEHSFPFQFLLPATAPTSFEGPFGKIVHQ<br>VRAAIHTPRFSKDHKCSLVFYILSPLNLNSIPDIEQPNVASATKKFSYKLVKTGSVVLTASTDLRG<br>YVVGQALQLHADVENQSGKDTSPVVASLLQKVSYKAKRWIHDVRTIAEVEGAGVKAWRRA<br>QWHEQILVPALPQSALPGCSLIHIDYYLQVSLKAPEATVTLPVFIGNIAVNHAPVSPRPGLGLP<br>PGAPPLVVPSAPPQEEAEAEAAAGGPHFLDPVFLSTKSHSQRQPLLATLSSVPGAPEPCPQD<br>GSPASHPLHPPLCISTGATVPYFAEGSGGPVPTTSTLILPPEYSSWGYPYEAPPSYEQSCGGVE<br>PSLTPES* |
| Myristoyl-PhoCl-MCP | MGSSKSKPKDPSQRGGGGSSSSYPYDVPDYAVIPDYFKQSFPEGYSWERSMTYEDGGICIAT<br>NDITMEGDSFINKIHFKGTNFPPNGPVMQKRTVGWEASTEKMYERDGVLKGDVKMKLLLK<br>GGGHYRCDYRTTYKVKQKPVKLPDYHFVDHRIEILSHDKDYNKVKLYEHAVARNSTDSMDEL |

| | |
|---|---|
| | YKGGSGGMVSKGEETITSVIKPDMKNKLRMEGNVNGHAFVIEGEGSGKPFEGIQTIDLEVKEGAPLPFAYDILTTAFHYGNRVFTKYPRDYKDDDDKLYSQRGGGGSASNFTQFVLVDNGGTGDVTVAPSNFANGVAEWISSNSRSQAYKVTCSVRQSSAQNRKYTIKVEVPKGAWRSYLNMELTIPIFATNSDCELIVKAMQGLLKDGNPIPSAIAANSGIYAMASNFTQFVLVDNGGTGDVTVAPSNFANGVAEWISSNSRSQAYKVTCSVRQSSAQNRKYTIKVEVPKGAWRSYLNMELTIPIFATNSDCELIVKAMQGLLKDGNPIPSAIAANSGIY* |
| Free-MCP | MASNFTQFVLVDNGGTGDVTVAPSNFANGVAEWISSNSRSQAYKVTCSVRQSSAQNRKYTIKVEVPKGAWRSYLNMELTIPIFATNSDCELIVKAMQGLLKDGNPIPSAIAANSGIYAMASNFTQFVLVDNGGTGDVTVAPSNFANGVAEWISSNSRSQAYKVTCSVRQSSAQNRKYTIKVEVPKGAWRSYLNMELTIPIFATNSDCELIVKAMQGLLKDGNPIPSAIAANSGIYGSS* |

Legend: MCP, Linker, Photocleavable domain (PhoCl), EV-targeting moiety, HA-tag, FLAG-tag

**Supplementary Table 4: DNA plasmid overview**

| Name | Function | Source |
|---|---|---|
| pMD2.G | VSV-G expression | Addgene #12259 |
| PSPAX2 | 2nd generation lentiviral packaging plasmid | Addgene #12660 |
| lentiCas9-Blast | Transient/stable expression of spCas9, labeled with a Flag-tag | Addgene #52962 |
| pCMV-ABE7.10 | Transient expression of adenine base editor ABE7.10 | Addgene #102919 |
| ABE8e | Transient expression of adenine base editor ABE8e | Addgene #138489 |
| Abe8e-dimer | Transient expression of adenine base editor ABE8e-dimer, containing an additional ecTadA domain | Addgene #138490 |
| Sp-dCas9-VPR | Transient expression of transcriptional activatory dCas9-VPR | Addgene #63798 |
| lentiGuide-puro | Transient/stable expression of WT sgRNAs | Addgene #52963 |
| lentiGuide-stoplight-puro | Transient/stable expression of WT sgRNA targeting the stoplight reporter construct | Previously published[34] |
| lentiGuide-TRE-puro | Transient/stable expression of WT sgRNA targeting the Tet Responsive Element | Previously published[34] |
| Lentiguide-ABE stoplight-puro | Transient/stable expression of WT sgRNA targeting the adenine base editor stoplight reporter construct | Generated for this study |
| lenti-sgRNA(MS2)-zeo | Transient/stable expression of MS2-2.0 sgRNAs | Addgene #61427 |
| lenti-Stoplight-sgRNA(MS2)-zeo | Transient/stable expression of a MS2-2.0 sgRNA targeting the stoplight reporter construct | Generated for this study |
| lenti-TRE-sgRNA(MS2)-zeo | Transient/stable expression of a MS2-2.0 sgRNA targeting the Tet Responsive Element | Generated for this study |
| lenti-ABE Stoplight-sgRNA(MS2)-zeo | Transient/stable expression of a MS2-2.0 sgRNA targeting the adenine base editor stoplight reporter construct | Generated for this study |
| lenti-gRNA puro | Transient/stable expression of sgRNAs or crRNAs, no gRNA backbone present in backbone | Addgene #84752 |
| lenti-ABE Stoplight 1.1-gRNA puro | Transient/stable expression of a MS2-1.1 sgRNA targeting the adenine base editor stoplight reporter construct | Generated for this study |
| lenti-ABE Stoplight 1.2-gRNA puro | Transient/stable expression of a MS2-1.2 sgRNA targeting the adenine base editor stoplight reporter construct | Generated for this study |
| pInducer20 | 2nd generation lentiviral transfer plasmid for doxycycline-inducible protein expression | Addgene #44012 |

| | | |
|---|---|---|
| pDONR221-eGFP | Used to clone eGFP into pInducer20 using the Gateway LR Clonase II Enzyme Mix | Addgene #25899 |
| pInducer20-eGFP | Transient/stable expression of doxycycline / dCas9-VPR-inducible eGFP expression | Generated for this study |
| pHAGE2-EF1a-IRES-PuroR | 2nd generation lentiviral transfer plasmid for protein expression | Darrel Kotton Lab |
| pHAGE2-EF1a-IRES-NeoR | 2nd generation lentiviral transfer plasmid for protein expression | Darrel Kotton Lab |
| pHAGE2-CMV-Stoplight-IRES-NeoR | Transient/stable expression expression of the Cas9 stoplight reporter construct | Previously published[34] |
| pHAGE2-EF1a-ABE Stoplight-IRES-NeoR | Transient/stable expression of the adenine base editor / HDR stoplight reporter construct | Previously published[17] |
| pHAGE2-EF1a- MCP-CD63-IRES-PuroR | Transient/stable expression of tandem MCPs fused to CD63 | Generated for this study |
| pHAGE2-EF1a- MCP-PhoCl-CD63-IRES-PuroR | Transient/stable expression of tandem MCPs fused to CD63 with a photocleavable domain | Generated for this study |
| pHAGE2-EF1a- MCP-PhoCl-CD9-IRES-PuroR | Transient/stable expression of tandem MCPs fused to CD9 with a photocleavable domain | Generated for this study |
| pHAGE2-EF1a- MCP-PhoCl-CD81-IRES-PuroR | Transient/stable expression of tandem MCPs fused to CD81 with a photocleavable domain | Generated for this study |
| pHAGE2-EF1a- MCP-PhoCl-ARRDC1-IRES-PuroR | Transient/stable expression of tandem MCPs fused to ARRDC1 with a photocleavable domain | Generated for this study |
| pHAGE2-EF1a- myristoyl-PhoCl-MCP-IRES-PuroR | Transient/stable expression of a myristoylation tag fused to tandem MCPs with a photocleavable domain | Generated for this study |
| pHAGE2-EF1a-Free MCP-IRES-PuroR | Transient/stable expression of cytosolic tandem MCPs | Generated for this study |

**Supplementary Table 5: PCR primers**

| Target | Orientation | Sequence |
|---|---|---|
| GAPDH | Forward | 5’-ACAGTCAGCCGCATCTTC-3’ |
| GAPDH | Reverse | 5’-GCCCAATACGACCAAATCC-3’ |
| Stoplight MS2-2.0 sgRNA | Forward | 5’-CAGTACTCCGCTCGAGTGTT-3’ |
| TRE MS2-2.0 sgRNA | Forward | 5’-TCACTGATAGGGAGTTTTAGAGC-3’ |
| MS2-2.0 sgRNAs | Reverse | 5’-TGTTGGCCAAGTTGATAACGG-3’ |
